# Germ-cell specific eIF4E1B regulates maternal RNA translation to ensure zygotic genome activation

**DOI:** 10.1101/2022.07.27.501690

**Authors:** Guanghui Yang, Qiliang Xin, Iris Feng, Jurrien Dean

## Abstract

Translation of maternal mRNAs is detected before transcription of zygotic genes and is essential for mammalian embryo development. How certain maternal mRNAs are selected for translation instead of degradation and how this burst of translation affects zygotic genome activation remains unknown. Using gene-edited mice, we document that the eukaryotic translation initiation factor 4E family member 1B (eIF4E1B) is the regulator of maternal mRNA translation that ensures subsequent reprogramming of the zygotic genome. In oocytes, the germ-cell specific eIF4E1B binds to mRNAs encoding chromatin remodeling complexes as well as reprogramming factors to protect them from degradation and promote their translation in zygotes. These protein products establish an open chromatin landscape in one-cell zygotes and enable transcription. Our results define a program for rapid resetting of the zygotic epigenome that is regulated by maternal mRNA translation and provides new insight into the mammalian maternal-to-zygotic transition.

## Introduction

Terminally differentiated, transcriptionally quiescent mammalian gametes fuse at fertilization and must be reprogrammed to express embryonic genes^1^. Maternal products stored in oocytes direct modifications of the epigenome^2^ after which the embryonic genome orchestrates development^3^.Mechanisms controlling this maternal-to-zygotic transition are not fully understood. The earliest transcripts from mouse zygotic genes are detected in late 1-cell zygotes and are followed by a more extensive rise of gene expression in 2-cell embryos. The two waves of transcription are designated minor and major zygotic genome activation (ZGA), respectively^4^. Active translation in mammals occurs before activation of zygotic genes^5,6^ and mouse embryos arrest at the 1-cell stage if this translation is inhibited^7,8^. Why this early burst of translation is essential for embryogenesis remains unknown, but recent experiments suggest this early translation is highly selective^9,10^ as most maternal RNAs and proteins are rapidly cleared during the maternal-to-zygotic transition (MZT)^11,12^. Considering the brief temporal window between fertilization and the earliest zygotic gene transcription, we hypothesize that the maternal mRNA translation is highly regulated to ensure availability of factors for efficient zygotic gene reprogramming. Using a candidate gene approach and gene-edited mice, we identify an essential role for a germ-cell specific eukaryotic translation initiation factor 4E family member 1B (eIF4E1B) in maternal mRNA translation that is essential for the maternal-to-zygotic transition.

## Results

### Inhibition of maternal mRNA translation prohibits mouse zygotic development

To systematically confirm the effects of maternal mRNA translation on embryo development, we cultured *in vitro* fertilized mouse eggs in medium containing cycloheximide (CHX) or anisomycin to inhibit protein translation (Fig. 1a). Inhibition of protein synthesis was confirmed (Fig. 1b, c) and most embryos arrested at the 1-cell pronuclear stage (Fig. 1d). In agreement with previous reports, our results emphasize the importance of maternal RNA translation in ensuring embryogenesis^7,8^. To identify the possible regulator controlling maternal RNA translation, we selected multiple candidate genes and generated knockout mouse lines, from which we found *Eif4e1b* to be a regulator that further controls the maternal-to-zygotic transition and embryogenesis.

**Fig. 1.**
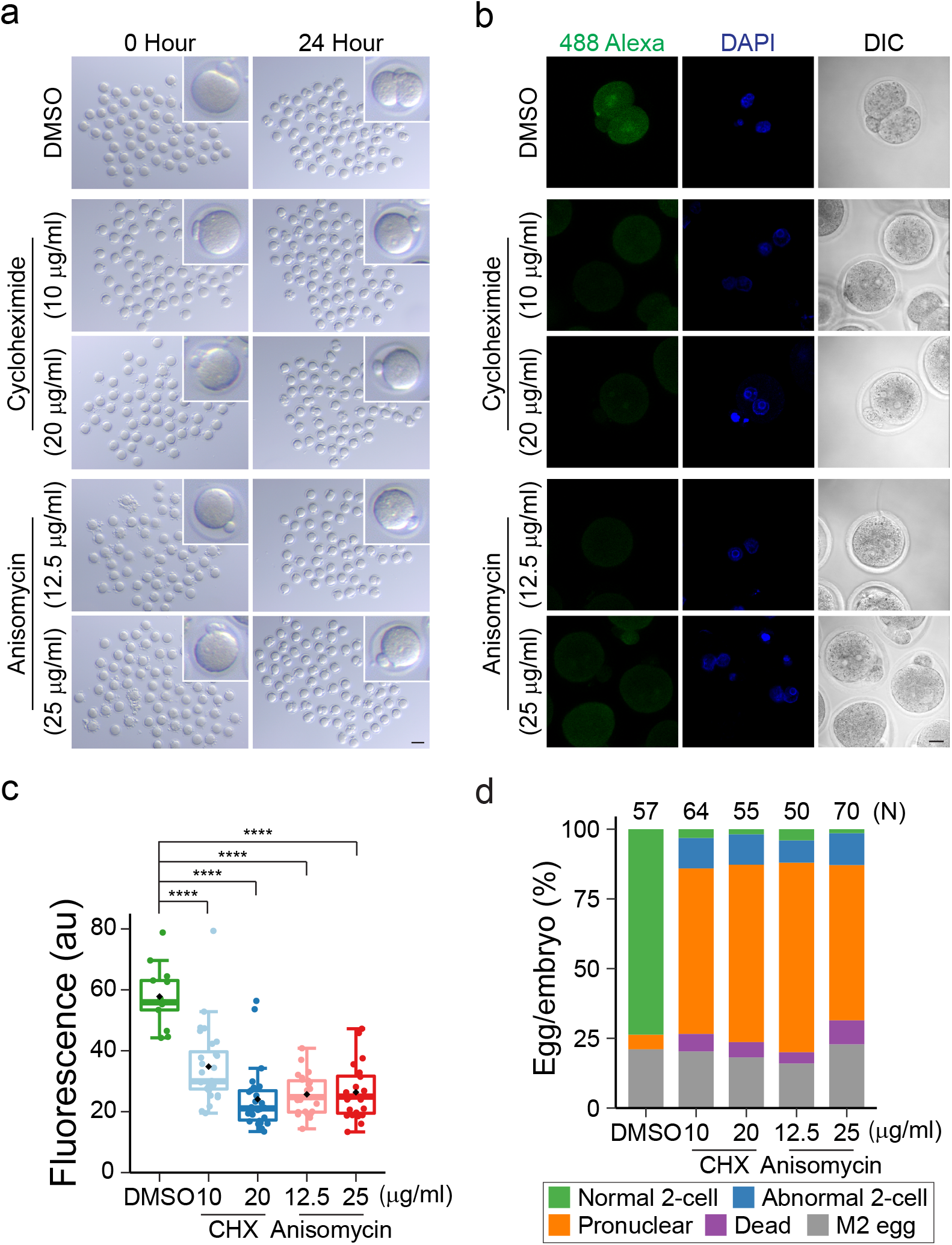
Inhibition of maternal mRNA translation prohibits mouse zygotic development. **A** Imaging M2 eggs before IVF (0 Hour) and embryos after IVF and drug treatment (24 Hour). Inset shows one enlarged representative egg/embryo (4.5× magnification). Scale bar, 100 μm. **b** Imaging nascent proteins 24 h after IVF. Representative images are shown. Inhibition of protein synthesis arrested embryos at the pronuclear stage while control embryos progressed to 2-cells 24 h after insemination. Scale bar, 20 μm. **c** Qualification of fluorescence signal in all embryos examined in **b**. The box plot includes the median (horizontal line) and data between the 25th and 75th percentile and each dot reflects the signal in one embryo. The black diamonds show average within each group. **** P < 0.0001, two-tailed t-test. **d** Ratio of embryos at different developmental stages with or without treatment 24 h after IVF. 4 h after insemination in IVF, embryos were washed and cultured in medium containing protein synthesis inhibitor cycloheximide (CHX) or anisomycin for additional 20 h. DMSO was used as the control.

### Maternal deletion of *Eif4e1b* arrests embryos at 2-cells

eIF4E1B is a member of the eIF4E (eukaryotic translation initiator factor 4E) super family that is essential for protein translation^13,14^. eIF4E1B shares 50% of its protein sequence with human eIF4E^15^, the founding member of the eIF4E family and, by analogy, binds to the 7-methylguanosine containing mRNA cap (Supplementary Fig. 1a). eIF4E1B protein from multiple species also share similar sequences^16^ (Fig. 2a), suggesting a conserved role. High level of *Eif4e1b* mRNA was detected in mouse ovary, with only trace expression in testis^17^ (Supplementary Fig. 1b). Using single embryo RNA-seq, we confirmed that *Eif4e1b* mRNA was abundant in mouse oocytes and persisted in 2-cell embryos (Fig. 2b, Supplementary Fig. 3a). Using a knock-in mouse line (*Eif4e1b*^*KI*^) in which FLAG and HA epitopes were added at the C terminus (Fig. 2c, Supplementary Fig. 2a, b), we also detected eIF4E1B protein in female germ cells (Supplementary Fig. 1c). eIF4E1B protein was detected in mouse oocytes and had increased expression in embryos until the late 2-cell stage as determined by immunostaining with samples derived from *Eif4e1b*^*KI*^ female mice (Fig. 2d). However, eIF4E1B protein was not detected in 4-cell embryos, agreeing with the absence of *Eif4e1b* expression after 4-cells^18^ (Supplementary Fig. 1d). Higher amount of *Eif4e1b* was detected in PN5 (pronuclear, stage 5) zygotes, comparing to that in metaphase II (M2) unfertilized eggs. Since (i) *Eif4e1b* is not detected in male germ cells in single cell RNA-seq experiments^19,20^; and (ii) zygotic gene transcription is low at the 1-cell stage where zygotic transcripts are poorly polyadenylated^21^, we speculate that the higher *Eif4e1b* abundance detected in PN5 zygotes is due to post-fertilization polyadenylation of maternal RNA^22^, which facilitated RNA capture in our poly(A) based single embryo RNA-seq experiment (Supplementary Fig. 3a). Taken together, the highly specific expression of eIF4E1B in oocytes made it an attractive candidate for translational control of maternal mRNA.

**Fig. 2.**
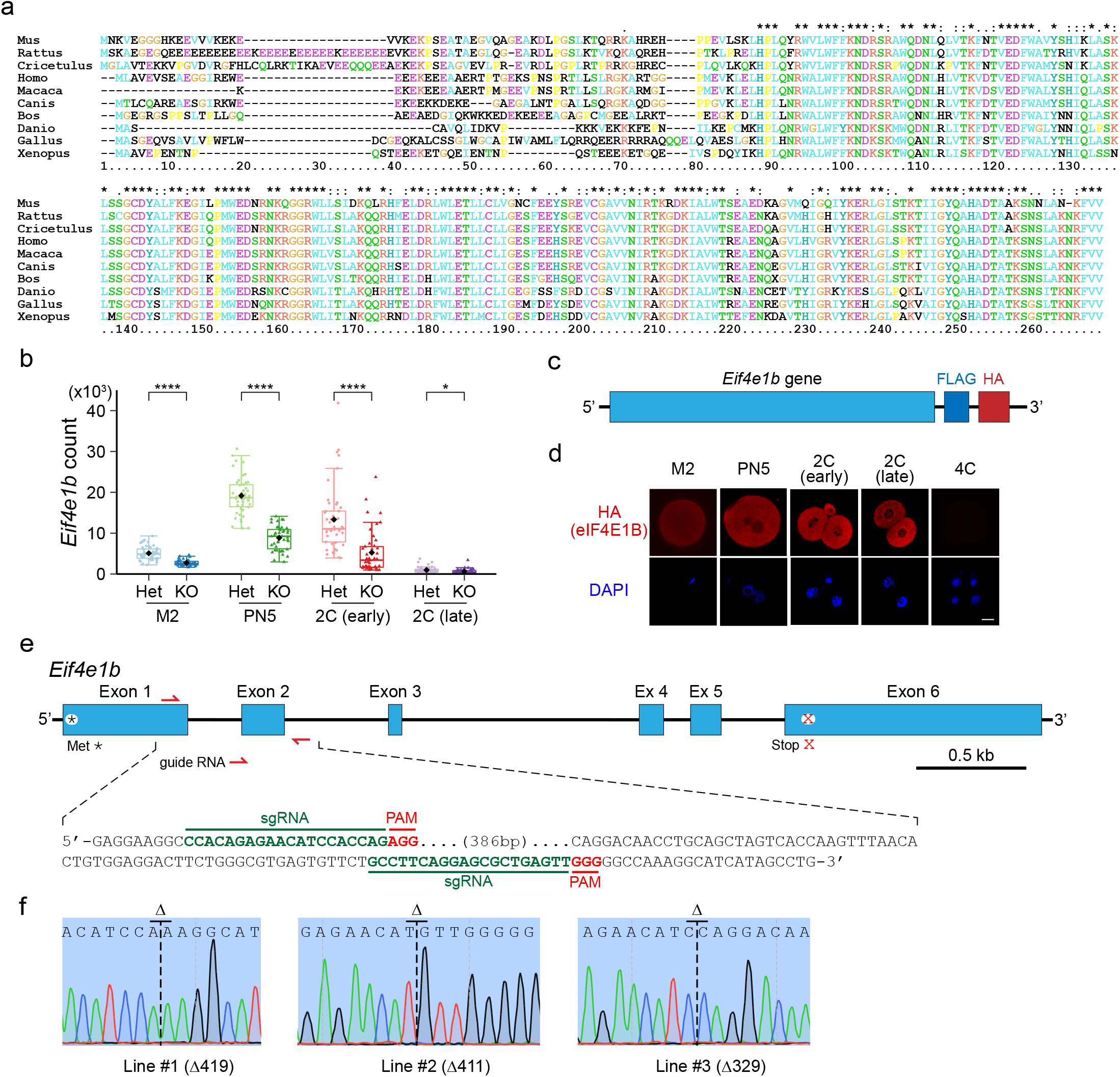
Generation of *Eif4e1b* gene-edited mouse lines. **A** Alignment of eIF4E1B protein sequences from multiple species. eIF4E1B sequences of *Mus musculus, Rattus norvegicus, Cricetulus griseus, Homo sapiens, Macaca mulatta, Canis lupus familiaris, Bos taurus, Danio rerio, Gallus gallus* and *Xenopus laevis* are aligned by ClustalX2. **b** Abundance of *Eif4e1b* mRNA in samples from *Eif4e1b*^*KO*^ and *Eif4e1b*^*Het*^ (control) female mice. All counts are normalized by ERCC spike-in. The box plot includes the median (horizontal line) and data between the 25th and 75th percentile and each dot reflects the result in one embryo. The black diamonds show average within each group. * P < 0.05, **** P < 0.0001, two-tailed t-test. **c** Schematic of the *Eif4e1b* gene locus in the *Eif4e1b*^*KI*^ mouse line with FLAG and HA tags at the C-terminus. **d** Immunofluorescence of eggs and embryos derived from *Eif4e1b*^*KI*^ female mice in which eIF4E1B has been fused with FLAG and HA tags at the C-terminus. Anti-HA antibody was used to visualize the eIF4E1B fusion protein. DAPI was used to visualize the nuclei. Scale bar, 20 μm. **e** Schematic of the *Eif4e1b* gene (upper) and sequences of sgRNAs (lower) for generation of *Eif4e1b*^*KO*^ mouse lines. *, initiator methionine; x, stop codon. **f** Sanger DNA sequencing at the *Eif4e1b* gene locus of the 3 knockout mouse lines.

To explore its function, we also generated *Eif4e1b* null mice using CRISPR/Cas9. After confirmation by DNA sequence, three *Eif4e1b* knockout lines were obtained and designated Δ419, Δ411 and Δ329 (Fig. 2e, f and Supplementary Fig. 2c) according to the size of their deletion. Unless noted, subsequent experiments were performed with the Δ419 line which was designated *Eif4e1b*^*KO*^ for homozygous null and *Eif4e1b*^*Het*^ for heterozygous mice that were used as controls. Significant reduction of *Eif4e1b* mRNA in eggs and early embryos retrieved from *Eif4e1b*^*KO*^ females was confirmed by single embryo RNA-seq (Fig. 2b). Although the residual *Eif4e1b* transcripts in these eggs/embryos existed in multiple isoforms, all of them had lost the first two exons (Supplementary Fig. 2d) and were not able to produce functional eIF4E1B protein (Supplementary Fig. 2e). These results further confirmed that eIF4E1B function was completely abolished in the knockout line. Homozygous null mice from all *Eif4e1b* knockout strains grew to adulthood. Adult males had normal fertility (Supplementary Fig. 2f) and testis morphology (Supplementary Fig. 2g), but the female mice were infertile (Fig. 3a).

**Fig. 3.**
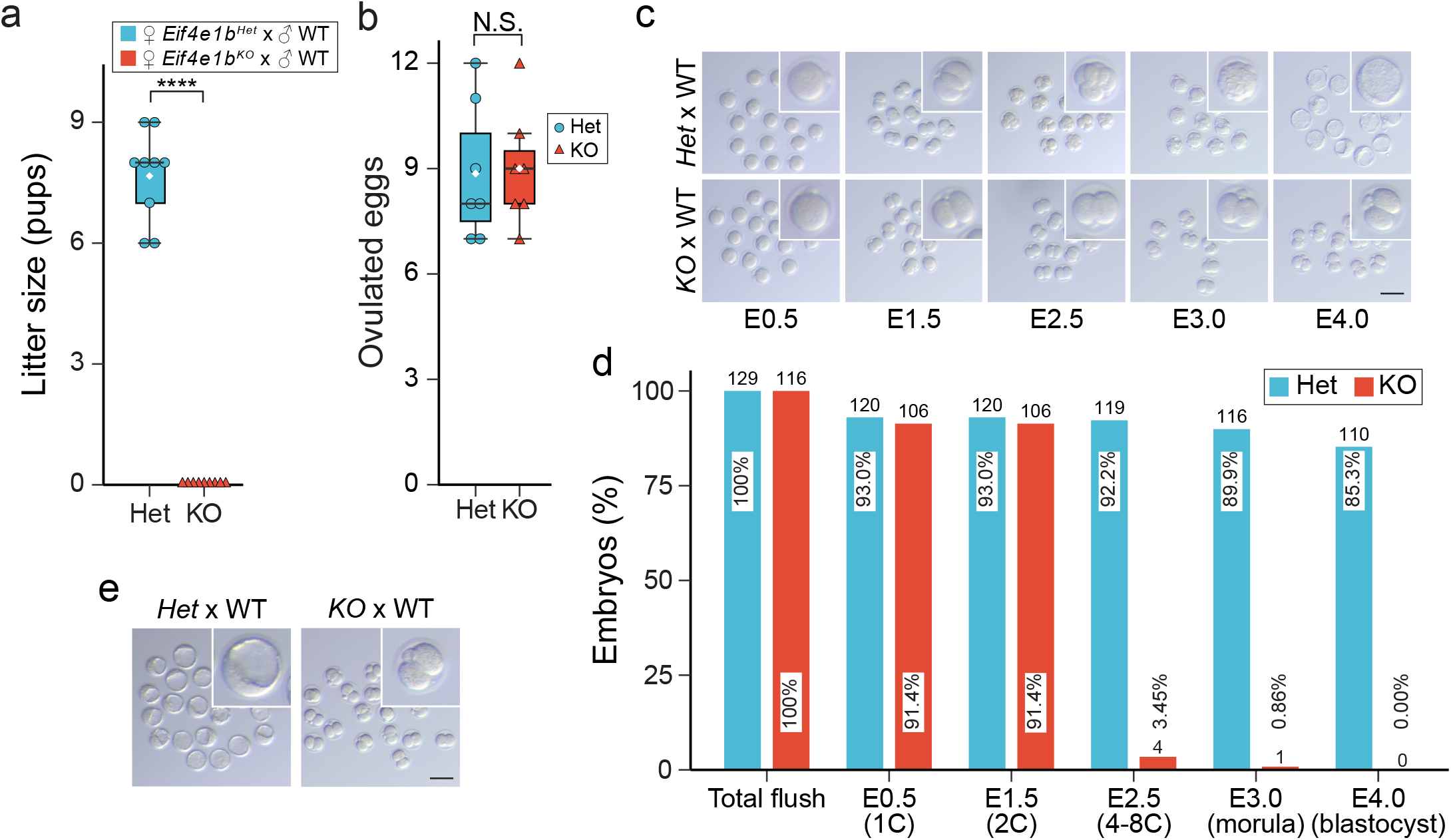
Maternal deletion of *Eif4e1b* leads to developmental arrest at 2-cells. **A** *Eif4e1b*^*Het*^ (control) and homozygous *Eif4e1b*^*KO*^ female litter sizes. **b** Number of ovulated eggs retrieved from *Eif4e1b*^*Het*^ or *Eif4e1b*^*KO*^ female mice after mating to WT males. The box plot includes the median (horizontal line) and data between the 25th and 75th percentile. Each dot or triangle reflects one observation. The white diamonds show average within each group. **** P < 0.0001, N.S. not significant, two-tailed t-test. **c** Representative images of *in vitro* cultured embryos from *Eif4e1b*^*Het*^ and *Eif4e1b*^*KO*^ females after mating with WT males at embryonic day 0.5 (E0.5), E1.5, E2.5, E3.0 and E4.0. Inset, 2.5× magnification. Scale bar, 100 μm. **d** Quantification of embryos as in **c**. Ratio of embryos at different stages is plotted. Total number of embryos is on top of each bar. **e** Images of embryos flushed from *Eif4e1b*^*Het*^ and *Eif4e1b*^*KO*^ female reproductive tracts at E3.5 after successful *in vivo* mating. Inset, 2.7× magnification. Scale bar, 100 μm.

*Eif4e1b*^*KO*^ female mice ovulate eggs (Fig. 3b) normally, which could be fertilized *in vitro* and *in vivo* but did not develop beyond 2-cell embryos. After mating control and *Eif4e1b*^*KO*^ females with wild-type (WT) males, fertilized zygotes were flushed from their oviducts and cultured *in vitro* for four days. The ratio of embryos that developed to different stages of pre-implantation development was determined (Fig. 3c, d). None of the embryos derived from *Eif4e1b*^*KO*^ female mice progressed beyond the 2-cell stage whereas control embryos became blastocysts (Fig. 3c). We confirmed that the 2-cell arrest occurred *in vivo*, by flushing control and *Eif4e1b*^*KO*^ female reproductive tracts at embryonic day 3.5 (E3.5) after mating with WT males (Fig. 3e). The arrested phenotype was also observed in the Δ411 and Δ329 lines (Supplementary Fig. 2h) which substantiated a role for *Eif4e1b* in developmental progression beyond 2-cells.

### Maternal ablation of *Eif4e1b* impairs ZGA

To investigate *Eif4e1b* function in early development, we adapted single-cell nucleosome, methylation, transcript sequencing (scNMT-seq)^23^ to single embryos (seNMT-seq) (Supplementary Fig. 3a). After mating hormonally stimulated control and *Eif4e1b*^*KO*^ female mice with WT males, zygotes were isolated for culture *in vitro*. Transcriptomes of these embryos, together with M2 eggs and PN5 zygotes, were analyzed using seNMT-seq (Supplementary Table 1). Most annotated protein-coding RNAs and long noncoding RNAs (lncRNAs) were detected in embryos from all stages which documented the efficiency of poly(A)-RNA capture and sequencing (Supplementary Fig. 3b). After quality control, principal component analysis (PCA) was performed to determine the relationship among samples (Fig. 4a). From the PCA plot, we calculated Euclidian distances between the centers of the two samples at each developmental stage to document differences between embryos from *Eif4e1b*^*KO*^ and control female mice (Supplementary Fig. 3c). Although 2-cell embryos within the same genotype exhibited significant heterogeneity (Fig. 4a), we consistently detected significant differences between 2-cell embryos derived from *Eif4e1b*^*KO*^ and control female mice (Fig. 4a, Supplementary Fig. 3c) which could account for the observed 2-cell arrest (Fig. 3c-e). In contrast, we did not see significant differences between the two genotypes in M2 eggs and PN5 zygotes (Fig. 4a, Supplementary Fig. 3c) which may reflect the absence of developmental delay at these stages (Fig. 3c). Interestingly, transcriptomes of M2 eggs and PN5 embryos with maternal *Eif4e1b* ablation were broadly down-regulated with few up-regulated transcripts (Supplementary Figs. 3d, e). This suggests accelerated mRNA clearance in these embryos and is consistent with the hypothesis that eIF4E1B binds and protects maternal mRNA from degradation.

**Fig. 4.**
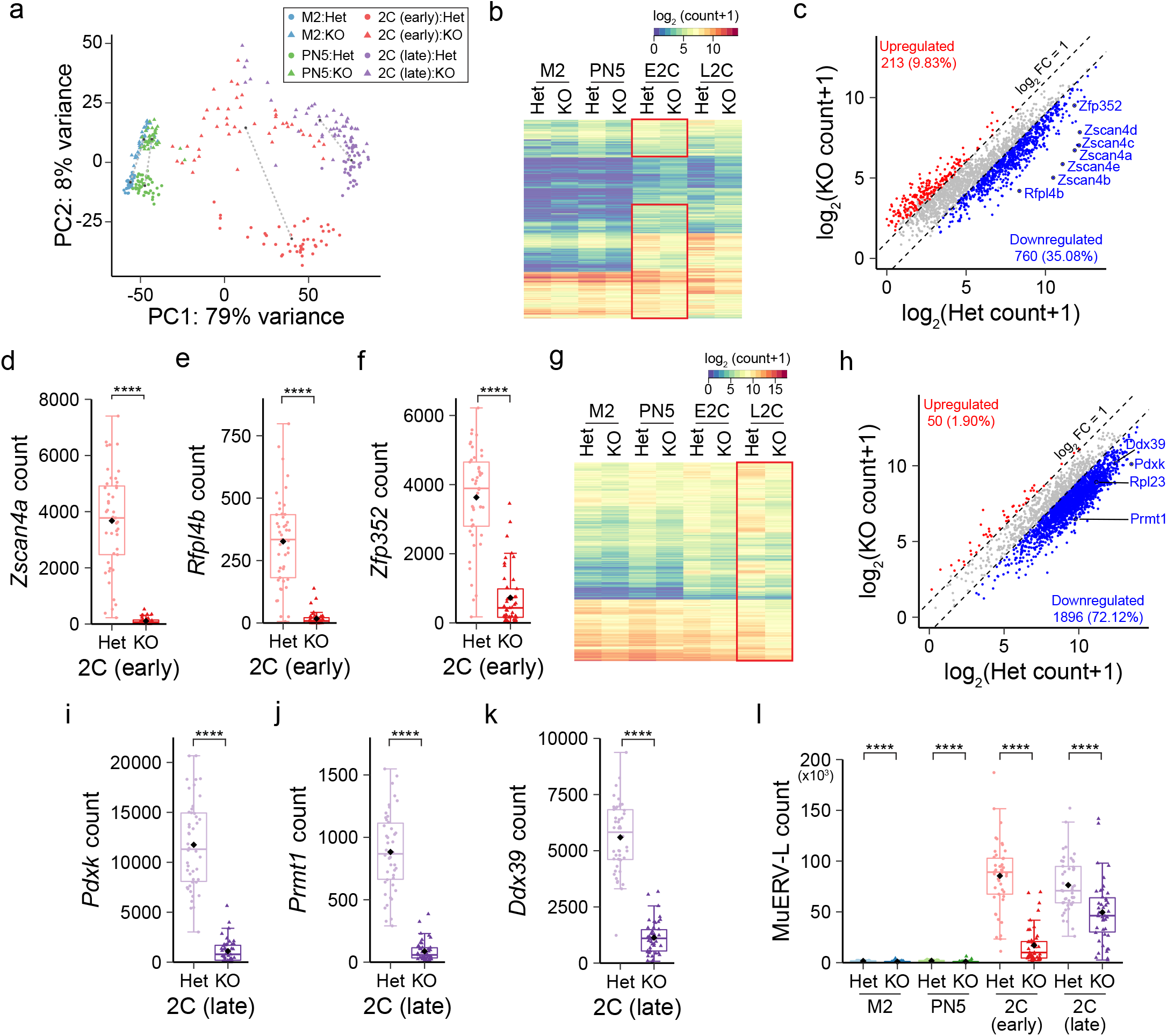
Maternal deletion of *Eif4e1b* impairs ZGA. **A** PCA plot of RNA-seq results of single embryos from *Eif4e1b*^*Het*^ (control) or *Eif4e1b*^*KO*^ female mice at different developmental stages. The length of dashed lines between cluster centers represents differences between samples. **b** Heatmap to show expression of all known minor ZGA genes at different stages. Note embryos from *Eif4e1b*^*KO*^ females have reduced expression of most minor ZGA genes at the early 2-cell stage (red box). **c**, Scatter plot documents differentially expressed RNAs expected to be transcribed during minor ZGA in early 2-cell embryos. Up-regulated and down-regulated RNAs are shown as red and blue dots, respectively. The total number of up-or down-regulated RNAs is labelled in each plot. mRNAs from multiple well-known minor ZGA genes are labeled in the plots. **d**-**f** Abundance of *Zscan4a, Rfpl4b* and *Zfp352*, three minor ZGA genes, at early 2-cell stage. **g** Heatmap to show expression of most known major ZGA genes at different stages. Note that embryos from *Eif4e1b*^*KO*^ females have reduced expression of almost all major ZGA genes at the late 2-cell stage as highlighted by the red box. **h**, Scatter plot documents differentially expressed RNAs expected to be transcribed during major ZGA in late 2-cell embryos. Up-regulated and down-regulated RNAs are shown as red and blue dots, respectively. The total number of up-or down-regulated RNAs is labelled in each plot. mRNAs from multiple well-known major ZGA genes are labeled in the plots. **i**-**k** Abundance of *Pdxk, Prmt1* and *Ddx39*, three major ZGA genes, at late 2-cell stage. **l** Abundance of transcripts from MuERV-L transposon in embryos from control or *Eif4e1b*^*KO*^ female mice at different developmental stages. All counts are normalized with ERCC spike-in. The box plot includes the median (horizontal line) and data between the 25th and 75th percentile and each dot reflects the count in one embryo. The black diamonds show average expression of the genes. **** P < 0.0001, two-tailed t-test.

The minor wave of mouse ZGA is detected in late 1-cell embryos about 14 hours after fertilization and continues into the early 2-cell stage^24^. Of the 2,166 reported minor ZGA transcripts^5,21,25^, 1,447 were down-regulated in embryos from *Eif4e1b*^*KO*^ female mice at the early 2-cell stage (Fig. 4b, Supplementary Table 2). Protein-coding RNAs normally up-regulated during minor ZGA^26^ (*e*.*g*., *Zscan4 cluster, Rfpl4b, Zfp352*) remained at low levels in early 2-cell embryos after maternal deletion of *Eif4e1b* (Fig. 4c-f and Supplementary Fig. 3f). Major ZGA follows the minor wave and these zygotic gene products direct subsequent development to establish the blueprint of early embryos^3^. Since the major ZGA is affected by the minor, it is not surprising that many major ZGA genes (*e*.*g*., *Prmt1, Pdxk, Ddx39*), including histone modifying enzymes were poorly expressed in late 2-cell embryos derived from *Eif4e1b*^*KO*^ female mice (Fig. 4g-k, and Supplementary Fig. 3g). Of 2,629 major ZGA transcripts^25^, 2,402 were downregulated in late 2-cell embryos derived from *Eif4e1b*^*KO*^ female mice (Fig. 4g, h, Supplementary Table 3), indicating near complete failure of major ZGA. These results suggest that maternal ablation of *Eif4e1b* causes both repression of genes that should be upregulated (Fig. 4b, g) as well as abnormally upregulated genes in late 2-cell embryos (Supplementary Fig. 3g). Both pathways affect ZGA and contribute to the 2-cell arrest.

Extensive activation of transposons in early mouse embryos has been reported and long terminal repeats (LTR) drive gene expression during ZGA^27,28^. MuERV-L has been used as a marker of successful zygotic genome activation^29^ and is reported to regulate LincGET as well as other pluripotency genes^30-32^. In embryos from *Eif4e1b*^*KO*^ females, MuERV-L was down-regulated at the 2-cell stage (Fig. 4l, Supplementary Table 4) as was global expression of LTRs (Supplementary Fig. 3h). *Dux* genes are reported to be among the earliest expressed zygotic genes in mice. Although originally thought to influence early embryo development^33-35^, their significance has been challenged more recently^36^. Only *Duxf4* expression was reduced in M2 eggs and PN5 zygotes from *Eif4e1b*^*KO*^ female mice (Supplementary Fig. 3i). Reduction of *Duxf3*, the most important *Dux* gene in mouse, was not observed, but higher levels were present in late 2-cell embryos from *Eif4e1b*^*KO*^ female mice (Supplementary Fig. 3j). A recent report suggests reduced *Duxf3* is necessary for embryo development beyond the 2-cell stage^37^. Thus. the altered *Duxf3* abundance after maternal *Eif4e1b* deletion may contribute to the observed 2-cell arrest but does not affect earlier embryo development. Taken together, our results suggest that maternal *Eif4e1b* deletion leads to systematical failure of the minor ZGA which causes major ZGA defects and leads to developmental arrest at the 2-cell stage.

### Maternal eIF4E1B reprograms zygotic chromatin accessibility

To further investigate mechanisms of impaired ZGA after maternal ablation of *Eif4e1b*, we exploited seNMT-seq to explore changes in DNA methylation and chromatin accessibility. The data for DNA methylation and chromatin accessibility from seNMT-seq are sparser in each single embryo compared to that from seRNA-seq. Thus, to overcome the difficulty from low sample size, we merged the results of all single embryos with the same genotype and from the same stage together to obtain a better global view of DNA methylation and chromatin accessibility. Although maternal ablation of *Eif4e1b* caused overall hyper-methylation of genomic DNA in early 2-cell embryos, no obvious changes were detected at the earlier PN5 stage (Fig. 5a, Supplementary Fig. 4a). DNA methylation at minor ZGA and major ZGA gene loci also showed no significant changes between PN5 zygotes from *Eif4e1b*^*KO*^ and control females and hyper-methylation at these regions was only detected at early 2-cell embryos derived from *Eif4e1b*^*KO*^ female mice (Fig. 5b and Supplementary Fig. 4b). We therefore conclude that rather than changes in genome DNA methylation, remodeling chromatin to render it more accessible provides the primary basis for early zygotic gene transcription. In this scenario, if maternal *Eif4e1b* is ablated, zygotic chromatin would remain inaccessible and lead to failed ZGA.

**Fig. 5.**
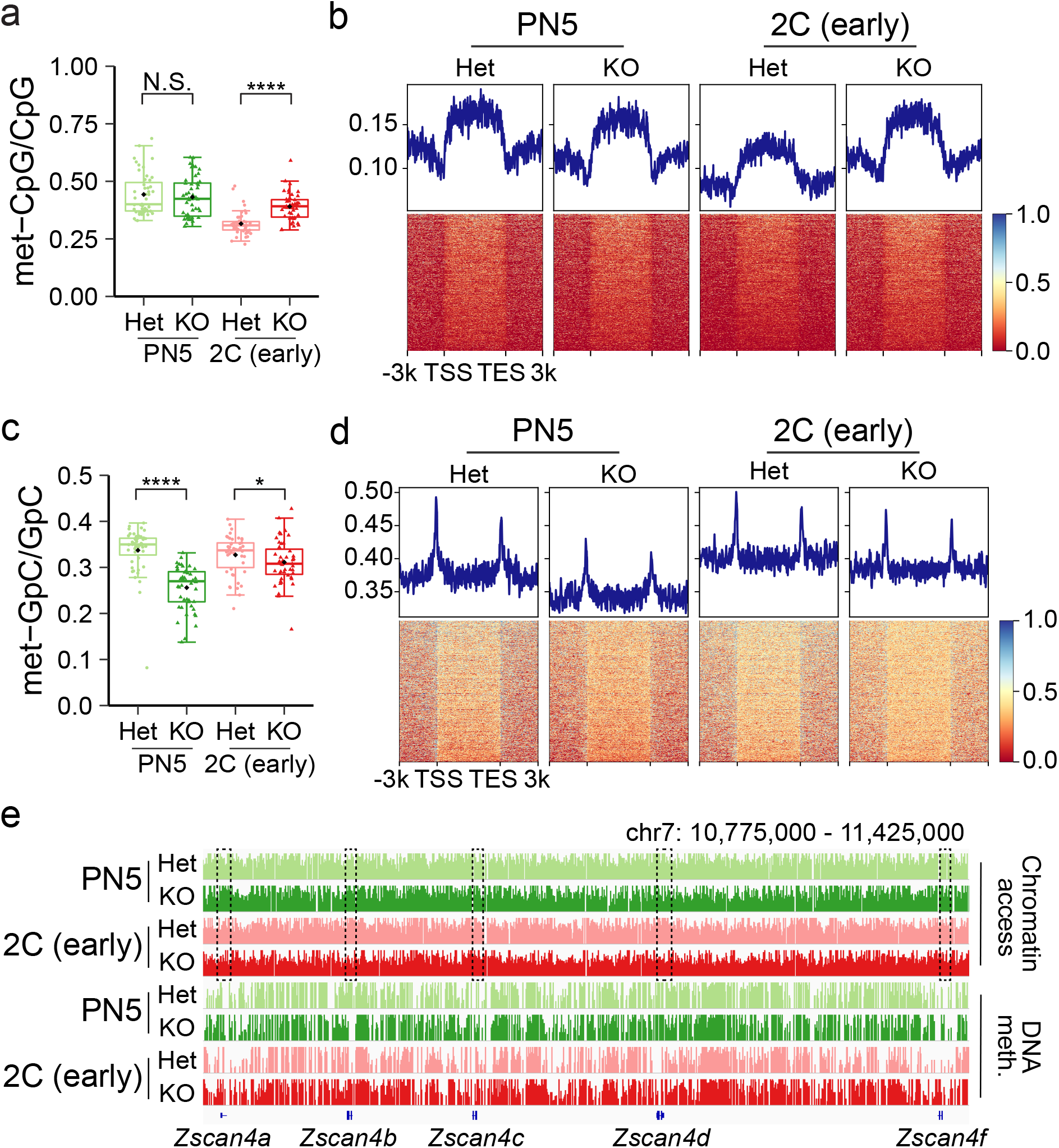
Maternal eIF4E1B reprograms zygotic chromatin accessibility. **A** Ratio of methylated CpG to document global DNA methylation. **b** DNA methylation profile at minor ZGA gene loci in PN5 zygotes and early 2-cell embryos from control and *Eif4e1b*^*KO*^ females. **c** Ratio of methylated GpC to show global chromatin accessibility. **d** Chromatin accessibility profile at minor ZGA gene loci in PN5 zygotes and early 2-cell embryos from control and *Eif4e1b*^*KO*^ females. **e** Integrated genomic view (IGV) to document chromatin accessibility and DNA methylation profiles at the *Zscan4* gene cluster. Note the lower chromatin accessibility at gene loci in embryos from *Eif4e1b*^*KO*^ females (framed). The box plot includes the median (horizontal line) and data between the 25th and 75th percentile. Each dot reflects the results from one embryo and the black diamonds show average within each group. N.S. not significant, * P < 0.1, **** P < 0.0001, two-tailed t-test.

Indeed, in contrast to the methylome changes, chromatin became less accessible in both PN5 zygotes and early 2-cell embryos in the absence of maternal eIF4E1B (Fig. 5c and Supplementary Fig. 4c). Severe and widespread decrease in chromatin accessibility at promoters of genes expected to express during the minor ZGA was observed in PN5 zygotes derived from *Eif4e1b*^*KO*^ female mice (Fig. 5d) including the *Zscan4* cluster (Fig. 5e). The lower chromatin accessibility continues until the early 2-cell stage, albeit to a lesser extent (Fig. 5d, e). The genomic locus of MuERV-L transposon also became less accessible (Supplementary Fig. 4d) after maternal ablation of *Eif4e1b*, consistent with the observed lower abundance of MuERV-L itself and downstream target transcripts. Reduced chromatin accessibility was also detected in early embryos derived from *Eif4e1b*^*KO*^ female mice at major ZGA gene loci, *e*.*g*., *Prmt1* (Supplementary Fig. 4e, f). These results support the hypothesis that maternal deletion of *Eif4e1b* fails to reset zygotic chromatin to an open structure which is the primary cause of failed ZGA (Fig. 4b, g).

### eIF4E1B binds mRNAs of chromatin remodeling complexes and reprogramming factors

As a member of the eukaryotic translation initiator factor 4E (eIF4E) family^13^, eIF4E1B binds the 7-methyguanosine cap of target mRNAs to promote translation and protect from degradation. In the absence of maternal eIF4E1B, target mRNAs are likely not efficiently translated and thus quickly degraded in M2 eggs and PN5 zygotes (Supplementary Fig. 3d, e). To confirm eIF4E1B binding and identify potential mRNA targets, we used M2 eggs and early 2-cell embryos from *Eif4e1b*^*KI*^ and control female mice (Fig. 2c, Supplementary Fig. 2a, b) to perform low-input RNA immunoprecipitation (RIP). There was no systematic difference in mapping input RNA to annotated genes (Fig. 6a and Supplementary Fig. 5a) from the two genotypes at the same developmental stage and immunoprecipitation (IP) results were compared without further normalization. eIF4E1B immunoprecipitated few annotated mRNAs in early 2-cell embryos derived from either control or *Eif4e1b*^*KI*^ female mice (Supplementary Fig. 5a, b) which suggested that eIF4E1B had little mRNA binding ability at this stage of development. In contrast, eIF4E1B bound more mRNAs (Supplementary Fig. 5a) transcribed from many fewer genes (Fig. 6b) in M2 eggs which is consistent with specific binding to a small subset of mRNAs in M2 eggs. In agreement with this result, we observed significant differences between control and *Eif4e1b*^*KI*^ samples of the RIP-seq data from M2 eggs (Supplementary Fig. 5b).

**Fig. 6.**
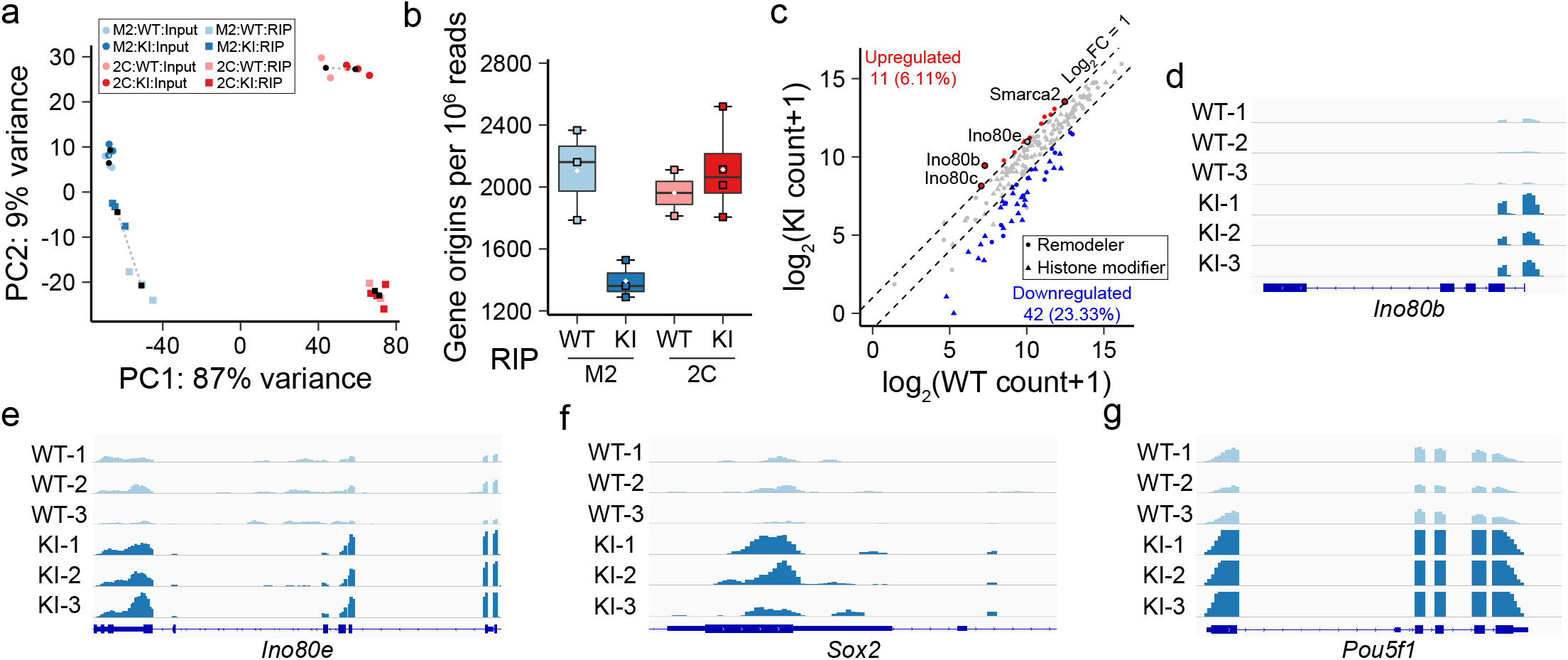
eIF4E1B binds to a subset of mRNAs in M2 eggs. **A** PCA analysis of input and immunoprecipitated transcripts after eIF4E1B-RIP. M2 eggs or early 2-cell embryos from *Eif4e1b*^*KI*^ female mice were used and WT eggs/embryos served as controls. **b** Gene origins per million reads from the RIP-seq data. Box plot includes the median (horizontal line) and data between the 25^th^ and 75^th^ percentile. Each small square reflects the results from one sample and the white diamonds indicate average in each group. **c** Scatter plot documents differentially expressed RNAs encoding known chromatin remodeling complex subunits and histone modifying enzymes as determined by the RIP-seq experiments using WT and *Eif4e1b*^*KI*^ M2 eggs. Up- and down-regulated RNAs are shown as red and blue dots, respectively. Several potential eIF4E1B mRNA targets are labeled. (**d**-**g**) Integrated genomic view (IGV) of eIF4E1B RIP-seq results at *Ino80b, Ino80e, Sox2* and *Pou5f1(Oct4)* loci in RIP-seq data from WT and *Eif4e1b*^*KI*^ M2 eggs.

RIP data in M2 eggs was reproducible within each genotype (Fig. 6a and Supplementary Fig. 5c) and we identified 3,436 RNAs that were more abundant in *Eif4e1b*^*KI*^ M2 eggs, representing candidate targets for eIF4E1B binding (Supplementary Fig. 5d, Supplementary Table 5). The RNAs underrepresented in the RIP-seq data from *Eif4e1b*^*KI*^ M2 eggs reflect non-specific immunoprecipitation observed in control M2 eggs (Supplementary Fig. 5d). Chromatin accessibility is regulated by remodeling complexes which can be affected by histone modifications^38^. We examined the RIP-seq results of the known 103 histone modifiers^39^ and 77 subunits of chromatin remodeling complexes^40^ in mouse to explore how eIF4E1B may affect the chromatin accessibility in early embryos (Fig. 6c, Supplementary Table 6). 10 of the 77 remodeling subunits showed significate upregulation in the RIP-seq results while only 1 of the 103 histone modifiers was upregulated. These results suggest eIF4E1B modulates chromatin accessibility by selective regulation of subunits of remodeling complexes. We focused on multiple members of the INO80 complex (Fig. 6d, e) and SMARCA2, a key member of the SWI/SNF complex (Supplementary Fig. 5e), which were potential eIF4E1B RNA targets. We also determined that *Sox2, Pou5f1* and *Polr1d* mRNA were additional potential eIF4E1B targets (Fig. 6f, g and Supplementary Fig. 5f, g). SOX2 and POU5F1 (OCT4) are well-known pluripotency factors that regulate early embryo development including zygotic genome activation^41-43^. These reprogramming factors interact with multiple remodeling complexes^44,45^ and may provide gene-specific localization during ZGA (Supplementary Fig. 6). POLR1D is an important component of RNA polymerase I, whose deletion leads to failed embryo development^46^. It is possible that POLR1D may facilitate translation of maternal or zygotic RNAs. By analyzing RNA sequences of potential eIF4E1B targets, we identified two motifs that may be used by eIF4E1B in selecting its targets (Supplementary Fig. 5h).

Taken together, our results suggest eIF4E1B can selectively bind mRNAs encoding chromatin remodeling proteins and reprogramming factors in oocytes to control zygotic chromatin accessibility through regulation of mRNA targets. The absence of binding to mRNAs encoding *Tet3* (Supplementary Fig. 5i) and other regulators of DNA methylation correlates with the absence of change in DNA methylation in zygotes derived from *Eif4e1b*^*KO*^ female mice. There was also an absence of binding to most mRNAs encoding histone modifiers (Fig. 6c, Supplementary Fig. 5j) and, thus, changes in chromatin accessibility appears to play the primary role in the ability of eIF4E1B to regulate ZGA.

### eIF4E1B promotes target mRNA expression

*Ino80b* knockout leads to embryonic lethality^47^ and genes bound by the INO80 chromatin remodeling complex have higher chromatin accessibility^48^. Continuous SWI/SNF activity is required for open chromatin structures^49,50^ and successful embryogenesis^51^. As reported, mRNA translation occurs in zygotes soon after fertilization^6^ and is essential for embryonic progression^7^. Considering the function of other members of the eIF4E family^13,14^, we were curious whether eIF4E1B reset zygotic chromatin accessibility by regulating protein translation of mRNA targets. The abundance of mRNA targets of eIF4E1B was decreased in M2 eggs and early embryos after maternal ablation of *Eif4e1b* (Fig. 7a). This was confirmed by expression of selected eIF4E1B mRNA targets during embryo development (Supplementary Fig. 7a-f) and is consistent with a protective effect on transcript stability by active translation^52^. Immunofluorescence using INO80B, IN80E and SMARCA2 specific antibodies determined their protein levels at different stages in early embryo development. Maternal ablation of *Eif4e1b* decreased stability of their mRNAs (Supplementary Fig. 7a-c) and cognate protein levels, especially in zygotes and early 2-cell embryos (Fig. 7b, c and Supplementary Fig. 7g, i, j). These observations were extended to reprogramming factors and confirmed by immunostaining (Fig. 7d, e and Supplementary Fig. 7h, k, l). This is consistent with the hypothesis that eIF4E1B binds essential mRNAs both to protect them from degradation and to promote their translation into proteins. Chromatin remodeling proteins translated could then modify the zygotic genome to create the open structures to facilitate ZGA.

**Fig. 7.**
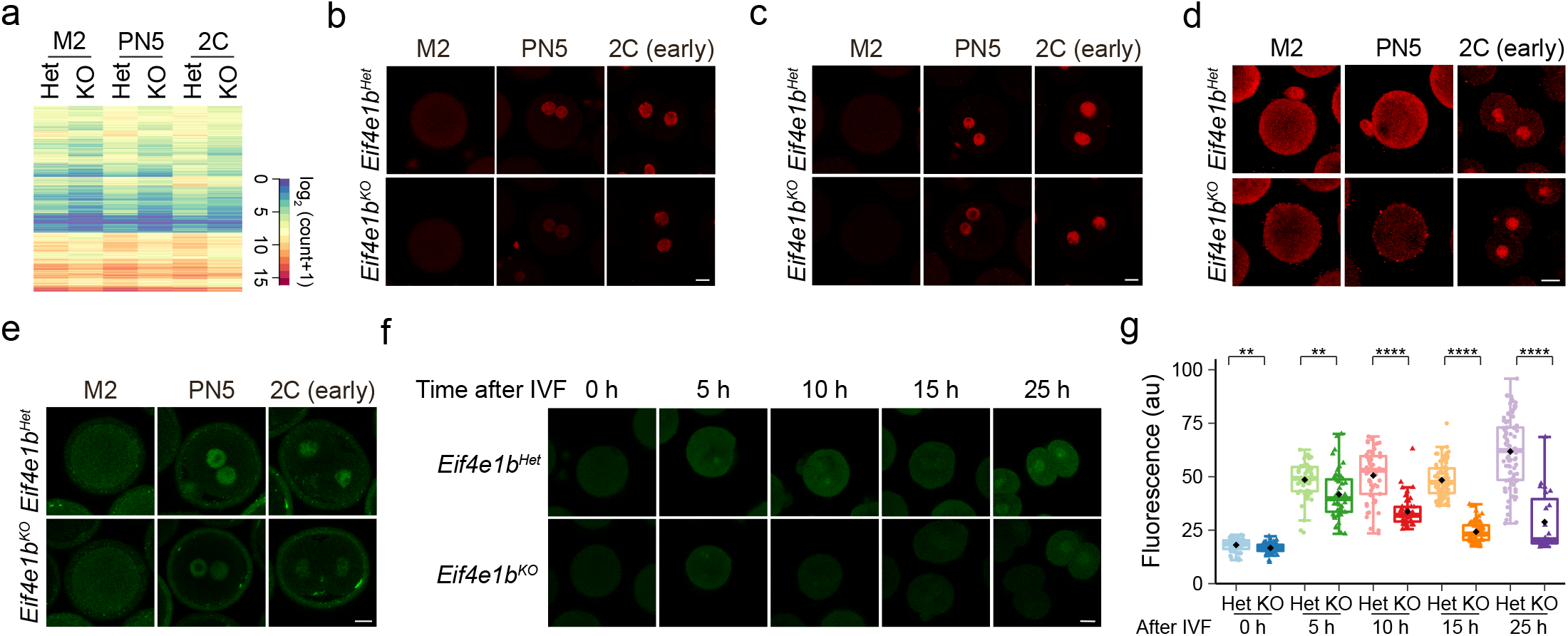
eIF4E1B controls translation of maternal mRNA in mouse zygotes. **A** Heatmap showing average expression of eIF4E1B RNA targets in embryos from *Eif4e1b*^*Het*^ and *Eif4e1b*^*KO*^ females at different developmental stages as determined by single embryo RNA-seq. All counts are normalized by ERCC spike-in. **b** INO80B protein expression in embryos from *Eif4e1b*^*Het*^ and *Eif4e1b*^*KO*^ females at different developmental stages. Scale bar, 20 μm. **c**-**e** Same as in b but for INO80E, SOX2 and OCT4 protein expression, respectively. The fluorescent signals are quantified in Supplementary Fig. 7**i**-**l. f** Imaging of nascent proteins in embryos derived from *Eif4e1b*^*Het*^ and *Eif4e1b*^*KO*^ females at different time points after IVF. The fluorescence signal was quantified in **g**. Scale bar, 20 μm. N.S. not significant, ** P < 0.01, **** P < 0.0001, two-tailed t-test.

### eIF4E1B controls maternal mRNA translation

To obtain a global view on protein expression controlled by eIF4E1B and to determine if maternal ablation of *Eif4e1b* affected protein synthesis in zygotes, we labeled nascent proteins in embryos after IVF and quantified signals at different time points. Significant reduction in protein biosynthesis was detected in zygotes and early 2-cell embryos from *Eif4e1b*^*KO*^ female mice (Fig. 7f, g). These results are consistent with eIF4E1B being essential for maternal mRNA translation in mouse zygotes.

## Discussion

After fertilization, the epigenome of mouse embryo must be reprogrammed to ensure transcription of zygotic genes^1,3^. Earlier investigations reported asymmetries in genomic DNA methylation in maternal and paternal pronuclei during reprogramming of the early zygotic epigenome. Similarly, differences of multiple histone modifications were observed between the paternal and maternal pronuclei in early zygotes^53^. However, later results indicated extensive demethylation of genome DNA occurred in both male and female pronuclei^54^ and that demethylation had little gene specificity. It was also noted that maternal and paternal alleles have similar chromatin accessibility at the late 1-cell (zygote) stage^55^, and embryonic gene expression has no significant parental allele preference^18^. These epigenomic data suggest that rapid reprogramming of zygotic chromatin accessibility may be the driver of minor and subsequent major ZGA. We now identify maternal eIF4E1B as a key, germ-cell specific component for translation of stored mRNAs in mouse 1-cell zygotes. eIF4E1B binds selectively to mRNAs encoding subunits of chromatin remodeling complexes and reprogramming factors in M2 eggs. We propose that after fertilization, but before ZGA, maternal eIF4E1B ensures translation of proteins required for resetting chromatin accessibility that enables expression of early zygotic genes (Fig. 8).

**Fig. 8.**
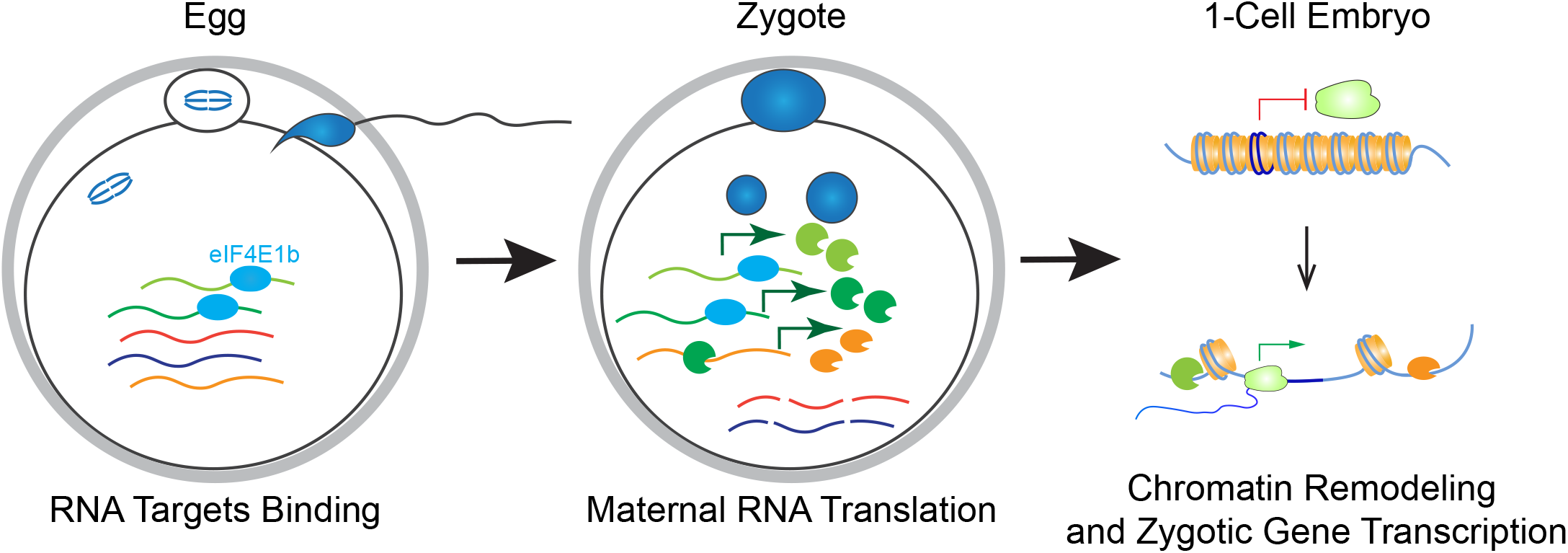
Working Model. eIF4E1B binds a subset of RNAs in M2 eggs. After fertilization, eIF4E1B bound mRNAs are rapidly translated into protein. Translation stabilizes the selected maternal mRNAs and prevents degradation. Their protein products remodel chromatin into a highly open state to enable transcription of the early zygotic genes that further establish early developmental programs. Maternal mRNAs and proteins are ultimately degraded during the maternal-to-zygotic transition.

Inhibition of maternal RNA translation arrests mouse embryos primarily at the 1-cell stage while embryos with maternal *Eif4e1b* ablation progressed to 2-cells. These results indicate that other factors participate in the regulation of maternal RNA translation. Our RIP-seq results suggest eIF4E1B has preference in binding RNA targets, but how targets are selected remains unkown. eIF4E1B is a relatively small protein with only one known domain and we suggest that additional co-factors may regulated target specificity for translation of maternal RNAs. Their identification will provide deeper insight into the maternal regulation of early embryogenesis.

Heretofore, investigations of the maternal-to-zygotic transition have focused on maternal product clearance and ZGA. Our current results document a program for rapid resetting of the early embryonic epigenome that is controlled by carefully orchestrated translation of maternal mRNA. A recent profiling of translated maternal mRNAs in mouse zygotes supports our findings of the importance of selective translation of chromatin remodeling complexes for ZGA^56^. Although the necessity of translation to trigger ZGA and the start of embryogenesis was previously suggested^6-8^, its regulatory mechanisms have remained unclear, and it has not been the focus of investigations into the maternal-to-zygotic transition. Our results confirm that maternal mRNAs are selectively regulated and explain why this burst of maternal mRNA translation is essential for embryo development. Our model supports the hypothesis that activation of early mouse embryogenesis is based on a genetic program pre-defined in female germ cells.

## Methods

### Ethics statement

All experiments with mice were conducted in accordance with guidelines of the National Institutes of Health under the Division of Intramural Research and NIDDK Animal Care and Use Committee approved animal study protocols (KO18-LCDB-18 and KO44-LCDB-19).

### Generation of CRISPR/Cas9 mutant mice

To establish the *Eif4e1b*^*KO*^ mutant mice, two CRISPR-Cas9 crRNA XT oligonucleotides^57^: 5′-CCACAGAGAACATCCACCAG -3′ and 5′-GCCTTCAGGAGCGCTGAGTT -3′ were synthesized by Integrated DNA Technologies. The crRNA was diluted (200 μM) in nuclease-free duplex buffer (Integrated DNA Technologies, Cat# 11010301). The two crRNA solutions were mixed with equal volumes of 200 μM tracrRNA (Integrated DNA Technologies, Cat# 1072533) separately and annealed into crRNA-tracrRNA duplexes using a thermocycler (Eppendorf). 1.5 μl of each crRNA-tracrRNA duplex solution was mixed with 1 μl S.p. HiFi Cas9 nuclease (Integrated DNA Technologies, Cat# 1081060) and 46 μl of advanced KSOM medium (Millipore, Cat# MR-101-D) to assemble the ribonucleoprotein (RNP) complex. The RNP was kept at room temperature for 10-30 min prior to use.

B6D2_F1_ (C57BL/6 ×DBA/2) female mice were hormonally stimulated with 5 IU of equine chorionic gonadotropin (eCG) followed 46-48 h later by 5 IU of human chorionic hormone (hCG) and then mated with B6D2_F1_ male mice. Zygotes in cumulus mass were released from the ampulla of the oviduct into M2 medium containing hyaluronidase (Millipore, Cat# MR-051-F) at embryonic day 0.5 (E0.5). Zygotes without cumulus were washed and transferred into advanced KSOM medium (Millipore, Cat# MR-101-D).

A NEPA21 electroporator (Nepa Gene) was used to deliver the RNP complex into zygotes (Poring pulse: voltage 225.0 V, pulse length 2.0 ms, pulse interval 50.0 ms, number of pulses 4, decay rate 10%, polarity +; Transfer pulse: voltage 20.0 V, pulse length 50.0 ms, pulse interval 50.0 ms, number of pulses 5, decay rate 40%, polarity +/-). 50 μl of RNP solution was aliquoted into the electrode (Nepa Gene, Cat# CUY505P5) along with 100 to 200 zygotes with a minimal volume of medium. The impedance of the solution was adjusted to ∼0.5 kΩ by changing the volume as determined by the NEPA21 electroporator. After electroporation, embryos were washed and cultured in advanced KSOM medium (37 °C, 5% CO_2_) for one additional day to obtain 2-cell embryos. Healthy 2-cell embryos were then transferred to the oviduct of pseudo-pregnant ICR females 1-day post coitus.

To establish a mouse line containing FLAG and HA tags fused at the C-terminus of *Eif4e1b*, crRNA XT was synthesized using the sequence 5’-CAACTTAGCAAACAAGTTTG-3’. RNP complexes containing 3 μl crRNA-tracrRNA duplex were assembled as described above. 12.5 μl of the RNP solution was mixed with 3 μl ssDNA (100 μM) in nuclease-free duplex buffer. Advanced KSOM was added to a final volume of 50 μl. Electroporation and embryo transfer were performed as described. The ssDNA for homologous repair^58^ was synthesized by Integrated DNA Technologies:

5’-CCAGAATCCACAGTGCAGTATAGTCTTCCTTGTCCATCAAGCAGCAAGATGAGGGTG CCCACTGAGTAGTGGCTGAAACCGGTCTCAGGCGTAGTCGGGCACGTCGTAGGGGTAGCTCCCTCCCTTATCGTCGTCATCCTTGTAATCACTGCCACCCACCACAAACTTGTTTG CTAAGTTGTTGCTCTTGGCAGCAGTGT-3’.

### Genotyping

Tail tips of mice were lysed in 200 μl of DirectPCR Lysis Reagent (Viagen Biotech, Cat# 102-T) with proteinase K (0.2 mg/ml, Sigma-Aldrich, Cat# 3115879001) at 55 °C for 4-16 h. To inactivate proteinase K, samples were incubated at 85 °C for 1 h. EmeraldAmp GT PCR Master Mix (Takara Bio USA, Cat# RR310A) and gene specific primers (Supplementary Table 7) were used to amplify specific DNA fragments. PCR was performed with an annealing temperature of 59 °C and 37 cycles using Mastercycler Pro (Eppendorf).

### Fertility assay

To test female fertility, pairs of *Eif4e1b*^*Het*^ (control) and *Eif4e1b*^*KO*^ female mice were harem mated with a WT male to determine the number and size of litters. *Eif4e1b*^*Het*^ and *Eif4e1b*^*KO*^ male mice were mated with WT females separately to determine male fertility.

### Histology and immunofluorescence

Mouse testes and ovaries were fixed in Bouin’s solution (Sigma-Aldrich, Cat# HT10132-1L) or 4% paraformaldehyde (PFA, Electron Microscopy Sciences, Cat# 15710) overnight at 4 °C for histology and immunostaining, respectively. Samples were embedded in paraffin, sectioned (5 μm) and mounted on slides prior to staining with periodic acid-Schiff (PAS) and hematoxylin.

For immunofluorescence, ovary sections were blocked with SuperBlock blocking buffer (ThermoFisher Scientific, Cat# 37515) containing 0.05% Tween-20 at room temperature for 1 h after de-waxing, rehydration, and antigen retrieval with 0.01% sodium citrate buffer (pH 6.0) (Sigma-Aldrich, Cat# C9999-100ML). The sections were then incubated with primary antibodies overnight at 4 °C. Goat anti-mouse antibody conjugated with Alexa Fluor 488 (1:500, Invitrogen, Cat# A-11001) or goat anti-rabbit antibody conjugated with Alexa Fluor 594 (1:500, Invitrogen, Cat# A-11012) were used to detect antigens and DNA was stained with DAPI in the mounting medium (ThermoFisher Scientific, Cat# P36941).

M2 eggs and embryos were fixed in 4% paraformaldehyde (PFA) for 30 min at room temperature and washed in phosphate-buffered saline (PBS, Invitrogen, Cat# 10010023) supplemented with 0.3 % polyvinylpyrrolidone (PVP, Sigma-Aldrich, Cat# PVP360-100G). Eggs/embryos were incubated in PBS with 0.3% BSA (Cell Signaling Technology, Cat# 9998S) and 0.1% Tween 20 (Sigma-Aldrich, Cat# P9416-50ML) for 2 h and stained overnight at 4 °C with anti-HA (Cell Signaling Technology, Cat# 3724S), anti-INO80B (Novus, Cat# NBP2-68903), anti-INO80E (Sigma, Cat# HPA043146), anti-SMARCA2 (Abcam, Cat# ab15597), anti-SOX2 (R&D Systems, Cat# MAB2018), anti-POLR1D (Proteintech, Cat# 12254-1-AP) or anti-OCT4 (Santa Cruz, Cat# sc-5279) primary antibodies. Goat anti-mouse or rabbit antibody conjugated with Alexa Fluor (Invitrogen) was used for immunofluorescent imaging. All the experiments were repeated at least three times and representative results from one replicate were presented.

### Single embryo NMT-seq

M2 eggs and embryos were collected from 6-8-week-old female mice. The females were injected intraperitoneally with eCG (5 IU) 46 h to 48 h prior to hCG (5 IU) injection and then co-caged with WT males. Fertilized zygotes were flushed from plugged females 16 h post hCG injection and cultured in M2 medium containing hyaluronidase to remove the cumulus mass. Zygotes without cumulus were then washed and cultured in advanced KSOM until sample collection. Embryos were collected at defined time points after hCG administration: PN5 (25 to 27 h), early 2-cell (35 h), late 2-cell (46 h). M2 eggs were collected 16 h post hCG injection without mating^59^. When collecting samples, M2 eggs or embryos were washed in PBS and transferred into acidic Tyrode’s solution (Millipore, Cat# MR-004-D) to remove zonae pellucidae. Single zona-free eggs/embryos were transferred into 8-well PCR strips containing 2.5 μl methyltransferase reaction mix which was comprised of 1 × M.CviPI Reaction buffer, 2 U M.CviPI (NEB, Cat# M0227S), 160 μM S-adenosylmethionine (NEB, Cat# B9003S), 1 U/μl RNasin (Promega, Cat# N2511), 0.1% IGEPAL CA630 (Sigma-Aldrich, Cat# I3021-50ML) in each well. The PCR strips were then incubated for 15 min at 37 °C in a thermocycler and the reaction was stopped by adding 5 μl RLT plus buffer (Qiagen, Cat# 1053393) to each well. The PCR strips with single eggs/embryos were frozen at -80 °C until library construction.

During RNA-seq library construction, 1 μl of pre-diluted (1:10^5^) ERCC spike-in was added to each well containing a single egg/embryo. RNA captured by the oligo-dT beads was converted into cDNA prior to amplification by 15 PCR cycles. After indexing, single embryo RNA-seq libraries from the same developmental stage were pooled together (usually 48 from *Eif4e1b*^*Het*^ and 48 from *Eif4e1b*^*KO*^ females) and purified with AMPure XP beads (Beckman, Cat# A63881) at a ratio of 1:0.6.

The supernatants containing genomic DNA after capture of RNA were processed following the scNMT-seq protocol^23^ with modified adapters:

First strand oligo: /5SpC3/TCGTCGGCAGCGTCAGATGTGTATAAGAGACAGNNNNNN Second strand oligo: GTCTCGTGGGCTCGGAGATGTGTATAAGAGACAGNNNNNN After PCR with Nextera XT dual indexing primers, single embryo DNA-seq libraries from the same developmental stage were pooled together and purified with AMPure XP beads at a ratio of 1:0.6. The quality of the pooled RNA-seq and DNA-seq libraries was confirmed by Bioanalyzer 2100 and each pooled library was sequenced (150 bp paired-end) in one lane on the Illumina HiSeq4000 platform (Novogene US).

### Low-input RNA immunoprecipitation (RIP)

200-250 M2 eggs or early 2-cell embryos were collected from WT or *Eif4e1b*^*KI*^ female mice and, after removing the zona pellucida, washed with PBS and transferred into 1.5 ml nuclease free centrifuge tubes with a minimal volume of PBS. The tubes were frozen immediately in dry ice and stored at -80 °C.

Low-input RNA immunoprecipitation was adapted by incorporating Smart-seq2^60^ and G&T-seq^61^ steps into the RIP-seq protocol^62^. Buffers from the EZ-Manga RIP kit (Millipore, Cat# 17-701) were used according to instructions from the manufacturer. 5 μl of anti-HA beads (ThermoFisher Scientific, Cat# 88836) was used for each RIP group and 20 μl of anti-HA beads (enough for 4 RIP groups) was prepared in one tube. The beads were separated on a magnetic rack, washed with 400 μl RIP wash buffer and resuspended in 1 ml RIP wash buffer (supplemented with 2% BSA) prior to rotation at 4 °C for 1 h to block non-specific binding. The blocked beads were washed on ice with 1 ml RIP wash buffer (supplemented with 2% BSA) and twice with 1 ml RIP wash buffer without BSA. The washed beads were resuspended in 800 μl RIP IP buffer supplemented with EDTA and RNase inhibitor.

100 μl freshly prepared lysis buffer from the EZ-Manga RIP kit containing protease inhibitor cocktail and RNase inhibitor was added to each previously frozen tube, tapped briefly, and kept on ice for 5 min. After freezing again on dry ice for 5 min, the thawed and lysed samples were used for the following experiments. 700 μl RIP IP buffer was added to each tube of lysate along with 200 μl of resuspended anti-HA beads. The tubes were rotated at 4 °C for 3 h and the beads were then separated magnetically. 200 μl supernatant was mixed with 360 μl RNAclean XP beads (1:1.8 ratio, Beckman, Cat# A63987) to purify the RNA. The RNAs bound by the RNAclean XP beads were used as input for each RIP group after washing (2X) with 80% ethanol. The remaining supernatant was discarded, and the beads were washed by 500 μl cold RIP wash buffer (6X) followed by the Smart-seq2 protocol to complete the library preparation: 18.2 μl elution buffer containing 9.2 μl RNase free water, 4 μl 10 μM oligo-dT30VN primer, 4 μl 10 μM dNTP mix and 1 μl RNase inhibitor (40 U/μl, Ambion, Cat# AM2682) was added to each tube containing RNAclean XP beads (input group) or anti-HA beads (IP group). The beads were triturated and transferred to individual wells of a PCR strip together with elution buffer. Elution was performed using a thermocycler with the following program: 55 °C 5 min, 70 °C 3 min. Other reagents used by Smart-seq2 for reverse transcription were mixed according to the volume of the elution buffer to a final volume of 21.8 μl. This reagent mix was added to each well of the PCR strip containing the eluted RNAs for reverse transcription. The cDNA in each well was then amplified following the Smart-seq2 protocol for 14 PCR cycles. The cDNA purification and tagmentation steps from the G&T-seq protocol were followed to provide indexing of the RIP libraries. Equal amounts of the RIP libraries (including the input) were mixed and sequenced (150 bp paired-end) in one lane on Illumina HiSeq4000 platform (Novogene US).

### Alignment of RNA-seq reads

The quality of FASTQ files was analyzed and confirmed by FastQC version 0.11.8. The reads were trimmed with Trimmomatic version 0.39 by indicating “NexteraPE-PE.fa” as the adapter sequence file^63^. The primary assembly of GRCm38 reference genome as well as the GTF annotation were downloaded from ENSEMBL (release 101). ERCC sequences as well as the corresponding GTF file were downloaded from the product page and concatenated to the end of the mouse reference genome and GTF files, respectively. The merged genome file and GTF file were used as references in downstream analysis. STAR version 2.7.6a was used to generate the genome indexes which were further used by STAR to align the trimmed FASTQ files^64^. Reads without pair-mates were also aligned by STAR and all the bam files from one sample were merged, sorted, and indexed by SAMtools version 1.12^65^. StringTie version 2.1.4 was used to generate counts of genes in the GTF reference^66^ which were further used for downstream analysis.

### Analysis of single embryo RNA-seq data

A total of 371 single embryo RNA-seq libraries were sequenced. Reads in each bam file that were aligned to the *Eif4e1b* deleted region as determined by *Eif4e1b*^*KO*^ genomic DNA were extracted and counted to confirm the genotype of each sample. Samples were deleted from downstream analyses if their genotypes were mislabeled or had high ratios of mitochondrial reads (more than 1.5 IQR above Q3). Samples with extremely high or low number of total reads (more than 1.5 IQR below Q1 or more than 1.5 IQR above Q3) were also considered outliers and deleted from downstream analysis. DESeq2 was used to analyze the cleaned RNA-seq data^67^ from 355 single embryos. The ERCC normalized gene count matrix was used for all plots. PCA and MA plots were generated using R. When Euclidian distances between different clusters were calculated, only PC1 and PC2 from the PCA plot were used. The percentage of variances from PC1 and PC2 was also considered during calculation. The gene biotype information was downloaded from Biomart^68^. Heatmaps illustrating RNA abundance detected in RNA-seq and the following experiments were generated by R heatmap.3 function with defined column order which represent different samples. The arrangement of the rows, which represent different RNAs, in the heatmaps was determined by the default arguments of heatmap.3 function.

### Analysis of transposable elements

The GTF file for mouse transposable element (TE) annotation was downloaded from the Hammell lab (Cold Spring Harbor) and ERCC spike-in GTF file was added to its end. The FASTQ reads were then re-aligned by STAR with this GTF file with the following parameter: “--winAnchorMultimapNmax 200 --outFilterMultimapNmax 100”. featureCounts was used to generate the expression table of annotated genes in the GTF^69^. The integer part of TE expression was used by DESeq2 and expression of ERCC spike-in was used to estimate the size factor for normalization.

### Alignment and processing of single-embryo DNA-seq data

Single-embryo DNA-seq data for analysis of DNA methylation and chromatin accessibility were aligned using HISAT-3N^70^ version 2.2.1-3n. Picard version 2.20.5 was used to remove duplicates in the bam files^71^. The methylated cytosines given by HISAT-3N were annotated by a home-made C++ program to identify CG and GC dinucleotides for analysis of DNA methylation and chromatin accessibility. Results of embryos from the same stage and of the same strain were merged and methylation rates of detected cytosines were calculated and transformed into bedGraph format. The bedGraph files were transformed in bigwig format and deepTools was used to generate the heatmaps covering genes that were interested^72^.

### Analysis of low-input RIP data

The number of reads that can be mapped to annotated genes as well as the total number of different genes that were mapped per million reads were calculated directly from bam files. The latter was used to check the gene origins of reads. Only RIP results from M2 eggs were used for downstream analysis as described in the text. The gene count matrix table was generated by Stringtie with the “-l 150” parameter and then used for the calculation and plotting. DESeq2, Biomart were used for analyses. Annotated transcripts with log_2_ fold change > 1 and padj value < 0.1 were considered differentially expressed and regarded as potential eIF4E1B targets. The bam files were first normalized with deepTools by FPKM and visualized with IGV^73^. Sequences of transcripts in the experimental group which had log_2_ fold change ≥ 2 and padj value ≤ 0.01 as determined by DESeq2 were analyzed by MEME-ChIP to identify shared motifs^74^.

### Re-analysis of ChIP-seq results

The bigwig files of the ChIP-seq experiments from GSE49137 and GSE87820 were used for analysis of INO80, SOX2, OCT4 recruitment in the genome. deepTools was used to generate the heatmap results with GTF annotation from ENSEMBL.

### Embryo treatment and imaging of protein synthesis

To determine effects of maternal mRNA translation on embryo development, M2 eggs were obtained from hormonally simulated WT females and incubated with sperm released from WT male epididymides for *in vitro* fertilization (IVF)^75^. 4 h later, unfertilized eggs and fertilized zygotes were washed and cultured in advanced KSOM medium supplemented with cycloheximide (CHX, Sigma-Aldrich, Cat# C7698-1G), anisomycin (Sigma-Aldrich, Cat# A9789-5MG) or DMSO (Sigma-Aldrich, Cat# D8418-50ML) as control for another 20 h before imaging. The Click-iT Plus OPP Alexa Fluor 488 Protein Synthesis Assay Kit (ThermoFisher Scientific, Cat# C10456) was used to determine nascent protein synthesis in each group of embryos. Nuclei were labeled with DAPI.

To quantify nascent protein synthesis in embryos fertilized from *Eif4e1b*^*Het*^ and *Eif4e1b*^*KO*^ eggs, IVF was performed as described. 4 h after insemination, the unfertilized eggs and fertilized zygotes in each group were washed and cultured in advanced KSOM medium. Zygotes were imaged 5 h, 10 h, 15 h, 25 h after insemination following the manufacturer’s instructions of the Click-iT Plus OPP Alexa Fluor 488 Protein Synthesis Assay Kit. M2 eggs were imaged before fertilization. All the experiments were repeated for at least three times and representative results from one replicate were presented.

### Quantification of fluorescence intensity

For all fluorescent staining experiments, the fluorescence intensity in each egg/embryo was quantified by ImageJ version 1.53k^76^ and then used for plotting in R.

## Acknowledgements

We thank all members of J.D. lab, especially Dr. Di Wu, for insightful comments.

## Author contributions

G.Y., Q.X. and J.D. conceived the project. G.Y., Q.X. and J.D. designed the experiments. G.Y., Q.X. and I.F. performed experiments. G.Y. analyzed results and wrote the manuscript with input from X.Q., J.D. revised the manuscript. All authors discussed and approved the manuscript.

## Data availability

The next generation sequencing data in this study has been deposited in the Gene Expression Omnibus website with accession number GSE180218. Source data for Fig. 1c, d; Fig. 3a, b; Fig. 7g; Supplementary Fig. 2f; Supplementary Fig. 7g-l are provided as a supplementary table (Supplementary Table 8). Other data are available in the main text and the supplementary materials. All other relevant data and materials that support the findings of this study are available from J.D. upon request.

