## Supplementary Figures 1 to 7 for "Germ-cell specific eIF4E1B regulates maternal RNA translation to ensure zygotic genome activation"

**This PDF file includes:**

Supplementary Figures 1 to 7

**Other Supplementary Materials for this manuscript include the following:**

Supplementary Tables 1 to 8

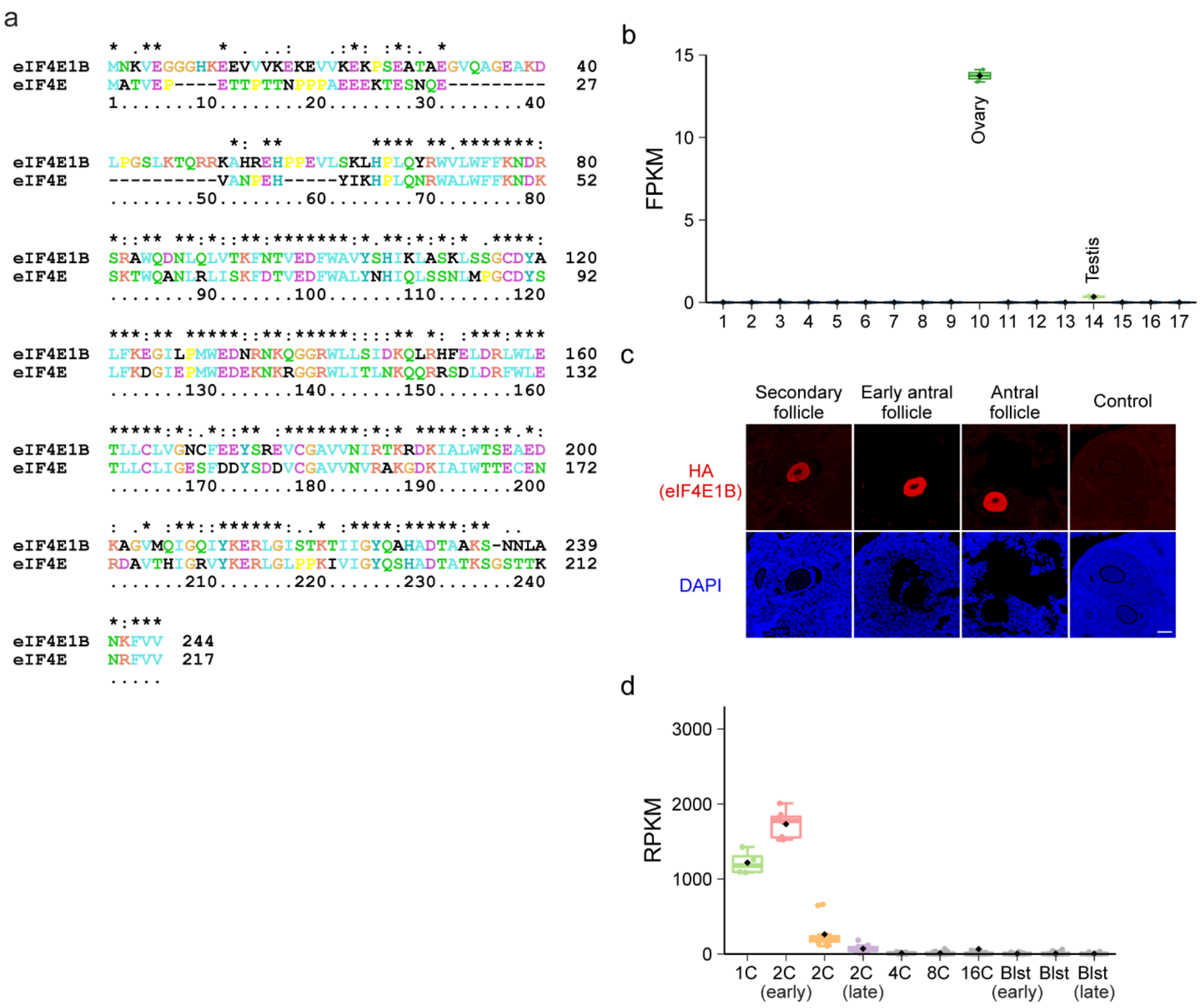


**Supplementary Figure 1. *Eif4e1b* expression in different samples. a** Alignment of eIF4E1B (NP001028441) protein sequence with that of eIF4E (NP031943). **b** Abundance (FPKM, fragments per kilobase of transcript per million mapped reads) of *Eif4e1b* transcripts in different tissues from published data[17]. 1, adrenal gland; 2, brain; 3, forestomach; 4, heart; 5, kidney; 6, liver; 7, large intestine; 8, lung; 9, muscle; 10, ovary; 11, small intestine; 12, spleen; 13, stomach; 14, testis; 15, thymus; 16, uterus; 17, vesicular gland. Box plot includes the median (horizontal line) and data between the $25^{th}$ and $75^{th}$ percentile. The black diamonds show average expression. **c** 6-week $Eif4e1b^{KI}$ mouse ovary sections (5 μm) were stained with anti-HA antibody to detect eIF4E1B-HA protein in follicles. DAPI was used to stain the DNA. Scale bar, 50 μm. **d**

Similar to **b**, but of mouse embryos from published data[18]. RPKM, reads per kilobase of transcript, per million mapped reads.

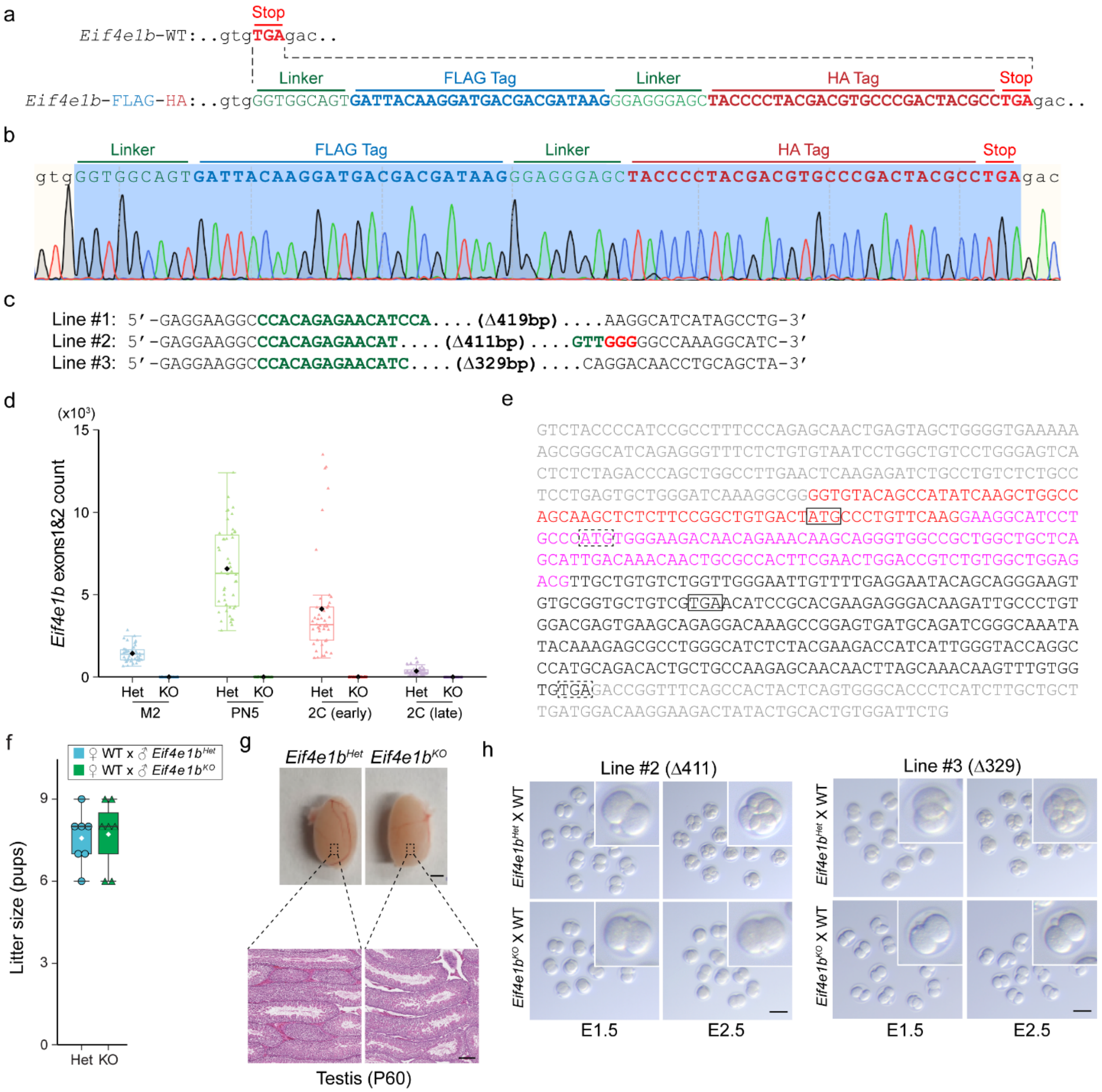


**Supplementary Figure 2. Characteristics of *Eif4e1b* gene-edited mouse lines. a** Design of the *Eif4e1b*$^{KI}$ mouse line with FLAG and HA tags fused to C-terminus of the eIF4E1B protein. **b** Sanger DNA sequencing at the *Eif4e1b*$^{KI}$ locus of the knock-in mouse line. **c** Genomic sequences at the *Eif4e1b* locus in 3 different *Eif4e1b*$^{KO}$ mouse lines with deletions ranging from 329 to 419 bp. **d** Abundance of reads overlap with *Eif4e1b* exon 1 or 2. The read counts are normalized by ERCC spike-in. Note that all residual *Eif4e1b* transcripts in samples from *Eif4e1b*$^{KO}$ female mice lose exons 1 and 2. **e** Sequence of residual *Eif4e1b* transcripts in eggs/embryos from *Eif4e1b*$^{KO}$

female mice. Sequence of the dominant isoform is shown. The 5’ and 3’ UTR sequences are shown gray. The 3rd and 4th exons of intact *Eif4e1b* are shared by all isoforms and are highlighted as red and magenta respectively. Two possible start codons and corresponding stop codons are framed. **f** *Eif4e1b*$^{Het}$ and *Eif4e1b*$^{KO}$ male litter sizes. The box plot includes the median (horizontal line) and data between the 25th and 75th percentile. The white diamonds show average within each group. **g** Morphology and histological sections of testes from *Eif4e1b*$^{Het}$ and *Eif4e1b*$^{KO}$ male mice. Scale bars, 2 mm (upper) and 50 μm (lower). **h** *In vitro* culture of embryos recovered from *Eif4e1b*$^{KO}$ line #2 (Δ411) and line #3 (Δ329) female mice after successful mating with WT males. Inset, 2.7× magnification. Scale bar, 100 μm.

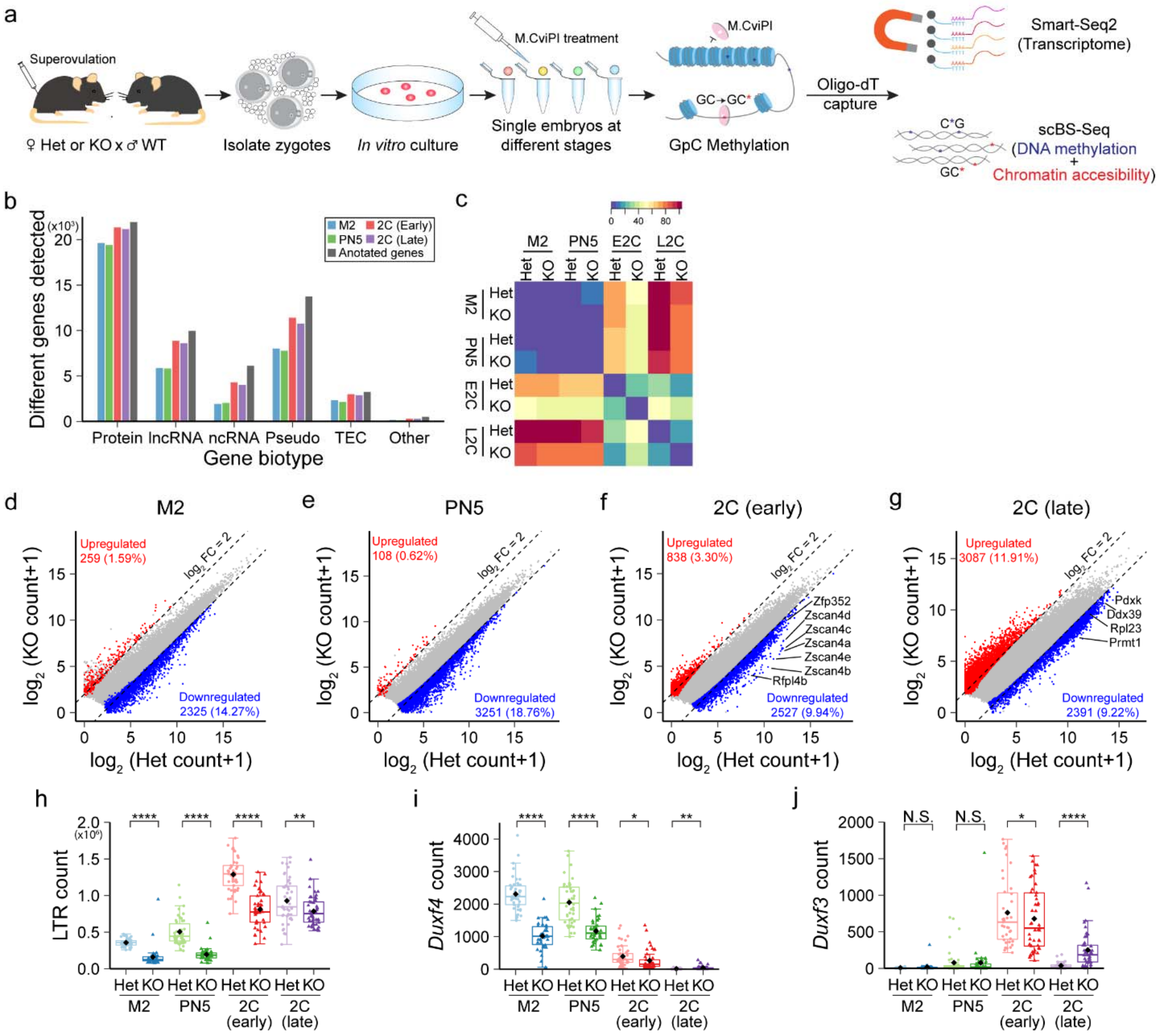


**Supplementary Figure 3. Analyses of single embryo RNA-seq data. a** Scheme of the single embryo NMT-seq. **b** Biotypes of all RNAs detected by single embryo RNA-seq. Note that most protein coding RNAs and lncRNAs are captured by the method. Protein, mRNA; lncRNA, long non-coding RNA; ncRNA, non-coding RNA; Pseudo, pseudogene RNA; TEC, to be experimentally confirmed. **c** Heatmap showing distances between different clusters of M2 eggs and early embryos in the PCA plot of Fig. 4**a**. Note the smaller differences in M2 and PN5 samples, even between samples from *Eif4e1b*$^{Het}$ and *Eif4e1b*$^{KO}$ females. (**d-g**) Scatter plots document differentially expressed RNAs in M2 eggs, PN5 zygotes and 2-cell (2C) embryos. Up-regulated and down-regulated RNAs are shown as red and blue dots, respectively. The total number of up- or down- regulated RNAs is labelled in each plot. mRNAs from multiple well-known ZGA genes are labeled in the plots of early 2-cell and late 2-cell embryos. **h** Abundance of transcripts from LTR transposons in embryos from control or *Eif4e1b*$^{KO}$ female mice at different developmental stages. (**i** and **j**) Abundance of *Duxf4* and *Duxf3* in embryos from control or *Eif4e1b*$^{KO}$ female mice at different developmental stages. All counts are normalized with ERCC spike-in. The box plot includes the median (horizontal line) and data between the 25th and 75th percentile and each dot reflects the count in one embryo. The black diamonds show average expression of the genes. N.S. not significant, * $P < 0.1$, ** $P < 0.01$, **** $P < 0.0001$, two-tailed t-test.

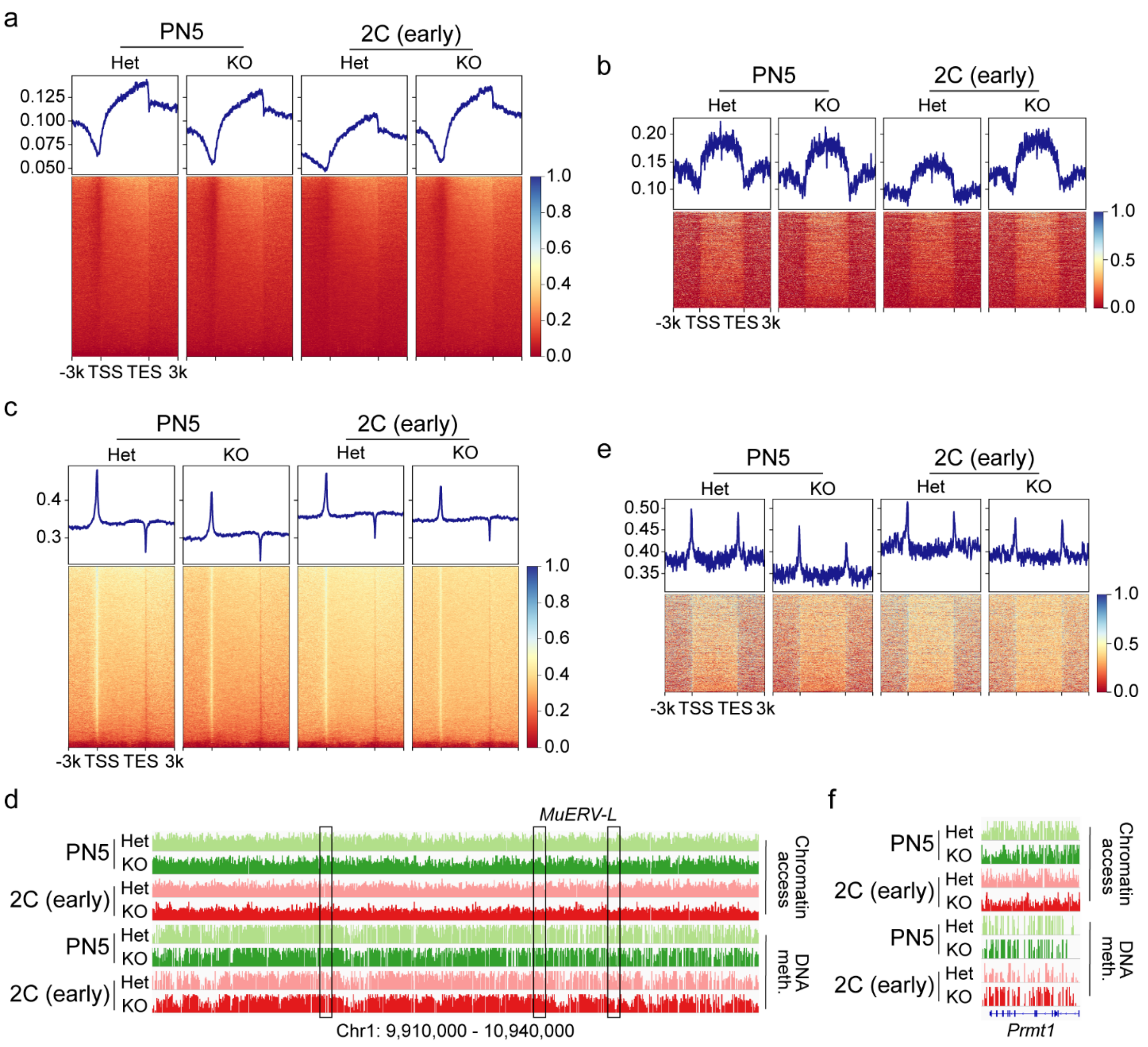


**Supplementary Figure 4. DNA methylation and chromatin accessibility in embryos from control and *Eif4e1b*$^{KO}$ females. a** DNA methylation at annotated genes to demonstrate global profiles in embryos from control and *Eif4e1b*$^{KO}$ female mice. **b** DNA methylation at major ZGA genes. **c** Chromatin accessibility at all annotated genes to show the global profile in embryos from control and *Eif4e1b*$^{KO}$ female mice. **d** Integrated genomic view (IGV) to show chromatin accessibility and DNA methylation profiles in chromosome 1 region containing the MuERV-L transposon. The regions coding MuERV-L are boxed. Note the lower chromatin accessibility around the boxed regions in embryos from *Eif4e1b*$^{KO}$ females. **e** Chromatin accessibility around major ZGA genes. **f** Similar to **d** but at the *Prmt1* gene locus.

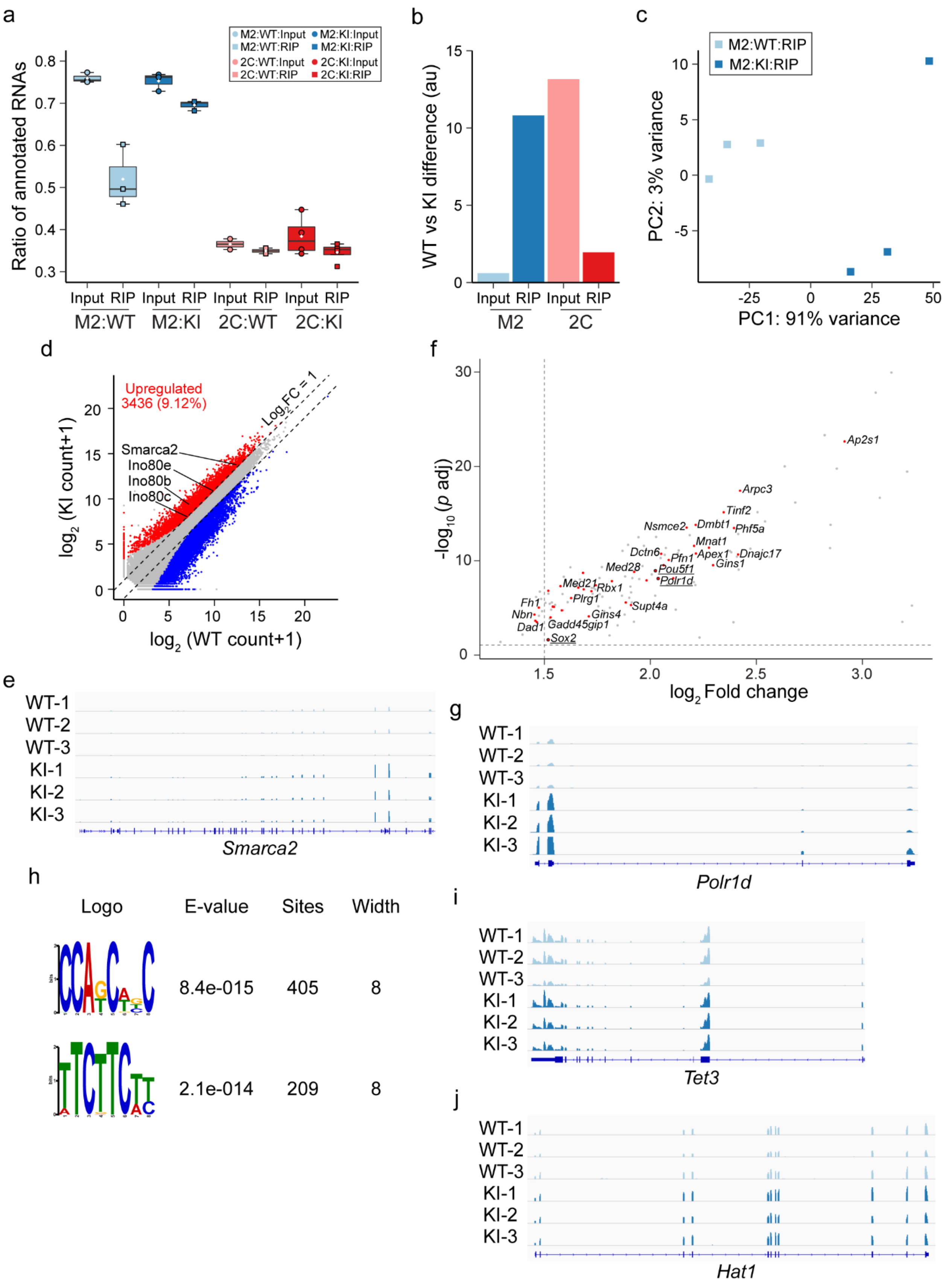

a
Ratio of annotated RNAs
M2:WT:Input
M2:WT:RIP
M2:KI:Input
M2:KI:RIP
2C:WT:Input
2C:WT:RIP
2C:KI:Input
2C:KI:RIP
Input RIP Input RIP Input RIP Input RIP
M2:WT M2:KI 2C:WT 2C:KI
b
WT vs KI difference (au)
Input RIP Input RIP
M2 2C
c
M2:WT:RIP
M2:KI:RIP
PC2: 3% variance
PC1: 91% variance
d
Upregulated
3436 (9.12%)
Smarca2
Ino80e
Ino80b
Ino80c
Log2FC = 1
log2 (KI count+1)
log2 (WT count+1)
f
Ap2s1
Arpc3
Tinf2
Nsmce2
Dmbt1
Phf5a
Mnat1
Dctn6
Apex1
Dnajc17
Pfn1
Gins1
Med28
Pou5f1
Pdlr1d
Med21
Rbx1
Plrg1
Supt4a
Fh1
Nbn
Gins4
Dad1
Gadd45gip1
Sox2
-log10 (p adj)
log2 Fold change
e
WT-1
WT-2
WT-3
KI-1
KI-2
KI-3
Smarca2
g
Polr1d
h
Logo E-value Sites Width
8.4e-015 405 8
2.1e-014 209 8
i
Tet3
j
Hat1


111

**Supplementary Figure 5. Analyses of RIP-seq data. a** Ratio of input and immunoprecipitated (RIP) reads that can be mapped to annotated mRNAs in the RIP-seq of M2 eggs and early 2-cell (2C) embryos from WT and *Eif4e1b*$^{KI}$ female mice. Box plot includes the median (horizontal line) and data between the 25$^{th}$ and 75$^{th}$ percentile. White diamonds indicate average in each group. **b** Length of the dashed lines in Fig. 6**a** showing differences between two group of samples. Variances of PC1 and PC2 are used for the calculation. **c** PCA analysis of input and RIP-seq data from M2 eggs. Note: The X axis captured almost all variance and embryos from the same genotype are close together while embryos from different genotypes are at a distance. **d** Scatter plot documents differentially expressed mRNAs found in the RIP-seq experiments of WT and *Eif4e1b*$^{KI}$ M2 eggs. Up- and down- regulated mRNAs are shown as red and blue dots, respectively. Several potential eIF4E1B mRNA targets are labeled. **e** Integrated genomic view (IGV) of eIF4E1B RIP-seq results at *Smarca2* gene locus in sequencing data from WT and *Eif4e1b*$^{KI}$ M2 eggs. **f** Volcano plot of transcripts expressed during embryogenesis and observed in RIP-seq. Each dot represents one transcript and those expressed during preimplantation development are red and labeled. *Sox2*, *Pou5f1*(*Oct4*) and *Polr1d* are also targets of eIF4E1B. **g** Integrated genomic view (IGV) of eIF4E1B RIP-seq results at *Polr1d* gene locus in sequencing data from WT and *Eif4e1b*$^{KI}$ M2 eggs. **h** RNA motifs shared by the transcripts detected in eIF4E1B RIP-seq. (**i** and **j**), Integrated genomic view (IGV) of eIF4E1B RIP-seq results at *Tet3* and *Hat1* gene loci in sequencing data from WT and *Eif4e1b*$^{KI}$ M2 eggs. No obvious binding of eIF4E1B is detected at these loci.

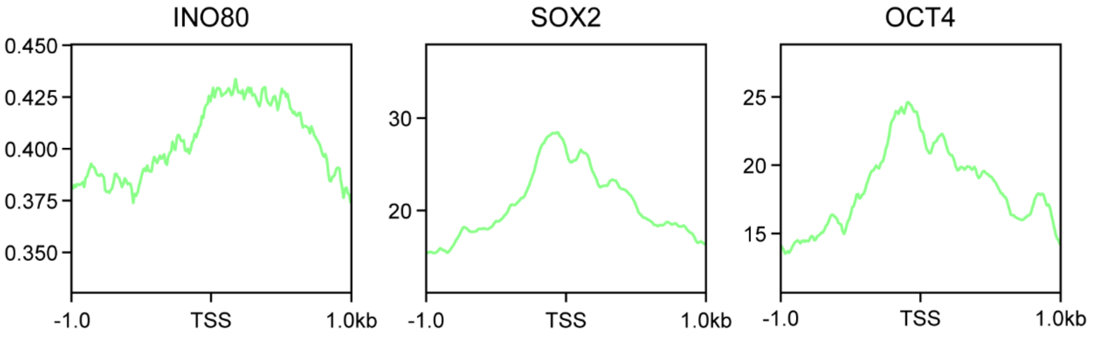


**Supplementary Figure 6. INO80, SOX2, OCT4 recruitment at minor ZGA gene promoters**. Preferences of INO80, SOX2, OCT4 recruitment in the genome were calculated from their ChIP-seq data from mouse embryonic stem cells. All of them show preferences in binding to minor ZGA gene promoters which may explain the specificity during remodeling of the zygotic chromatin.

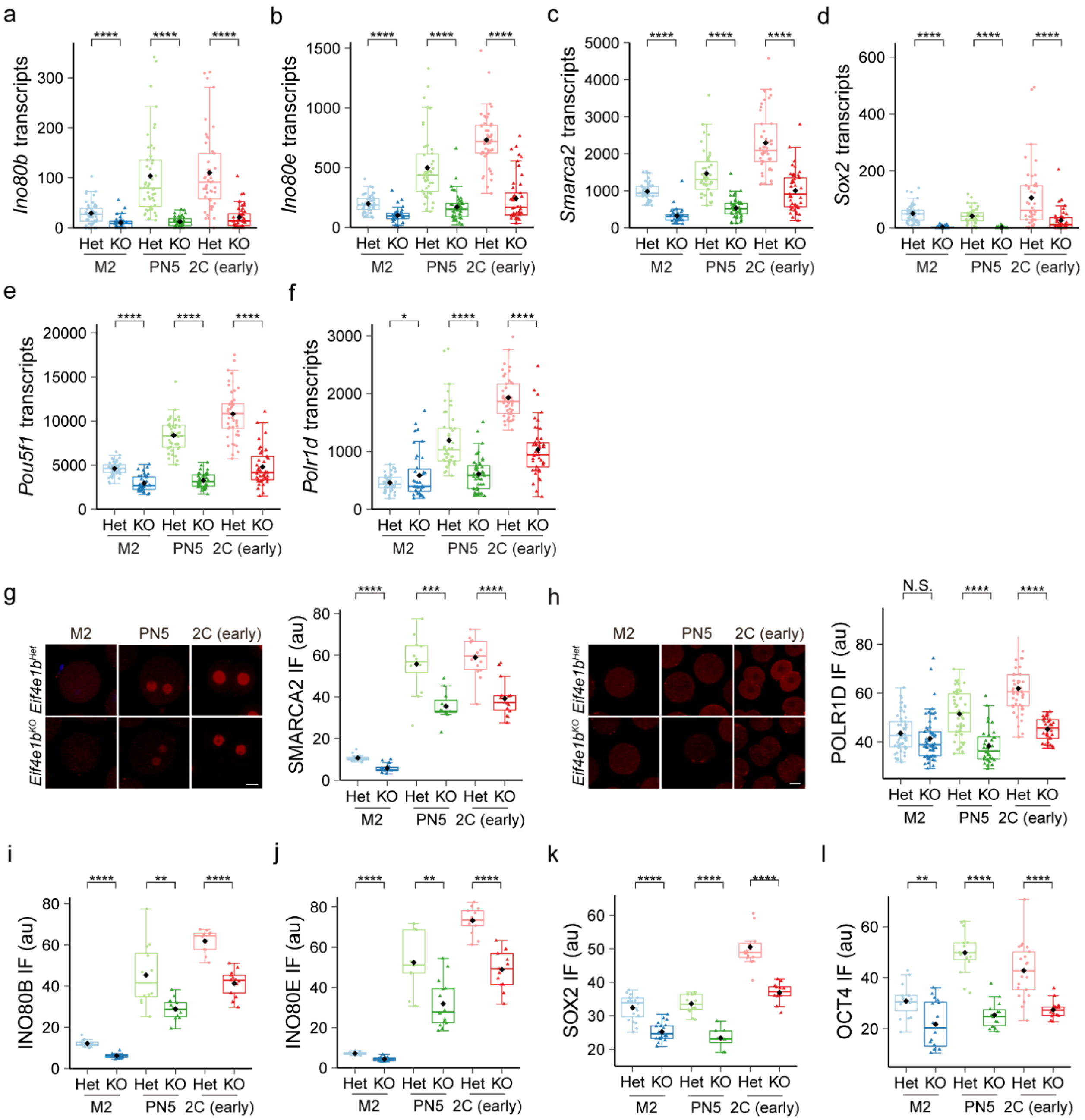


**Supplementary Figure 7. Expression of eIF4E1B targets in embryos from control and *Eif4e1b*$^{KO}$ females.** (**a-f**) Abundance of *Ino80b*, *Ino80e*, *Smarca2*, *Sox2*, *Pou5f1*(*Oct4*) and *Polr1d* in embryos from control or *Eif4e1b*$^{KO}$ female mice at different developmental stages as determined by single embryo RNA-seq. Note *Eif4e1b* maternal deletion leads to accelerated degradation of these RNAs due to lose of binding and protection. All counts are normalized with

ERCC spike-in and each dot or triangle reflects the abundance in one embryo. The black diamonds show average expression of the genes. The box plot includes the median (horizontal line) and data between the 25th and 75th percentile. * P <0.1, **** P < 0.0001, two-tailed t-test. **g** SMARCA2 protein expression in embryos from *Eif4e1b*$^{Het}$ and *Eif4e1b*$^{KO}$ females at different developmental stages. The fluorescent signals are quantified. **h** Same as in **g** but for POLR1D protein expression. Scale bar, 20 µm. (**i**-**l**) Quantifications of fluorescent signals in Fig. 7**b**-**e**. The box plot includes the median (horizontal line) and data between the 25th and 75th percentile. Each dot or triangle reflects the result in one embryo. The black diamonds show average in each group. N.S. not significant, ** P <0.01, **** P < 0.0001, two-tailed t-test.

**Supplementary Tables 1 to 8 are included in separate Excel (.xlsx) files.**

**Supplementary Table 1.** Normalized mean counts of all annotated genes at different stages of different strains.

**Supplementary Table 2.** Minor ZGA genes used for analysis.

**Supplementary Table 3.** Major ZGA genes used for analysis.

**Supplementary Table 4.** Normalized mean counts of all annotated transposable elements at different stages of different strains.

**Supplementary Table 5.** Differential gene expression analysis using M2 egg RIP-seq data.

**Supplementary Table 6.** Normalized counts of RNAs coding histone modifiers and subunits of chromatin remodeling complexes in RIP-seq using M2 eggs.

**Supplementary Table 7.** Primers and oligoes.

**Supplementary Table 8.** Source data for Fig. 1c, d; Fig. 3a, b; Fig. 7f; Supplementary Figures 2f, 8g-l.
